# Coevolution in a warming world: an experimental test of the geographic mosaic of coevolution

**DOI:** 10.1101/2025.11.06.686912

**Authors:** Quint Rusman, Tyler Figueira, Juan Traine, Florian P. Schiestl

## Abstract

According to the geographic mosaic theory of coevolution (GMTC), coevolution varies with abiotic and biotic environmental factors. We assessed this hypothesis using experimental plant-butterfly coevolution and by testing the effects of temperature and the presence of mutualistic bumblebees on co-divergence during six generations of selection. Butterflies are mutualistic by pollinating plants and antagonistic by ovipositing on plants from which caterpillars feed. We found unique plant-butterfly coevolutionary trajectories in response to abiotic and biotic factors: plants evolved strong herbivore resistance when exposed to either bumblebee presence or elevated temperatures, while their combination led to less strong plant-resistance evolution and the evolution of butterfly-foraging traits. We provide experimental proof for the GCMT and show rapid divergent coevolution to the combination of local abiotic and biotic conditions.

## Introduction

Coevolution is the reciprocal evolutionary change of ecologically interacting species driven by natural selection^1,2^, and one of the primary drivers of diversification across the tree of life^3,4^. According to the geographic mosaic theory of coevolution (GMTC), coevolution varies with abiotic and biotic environmental factors^3^. Environmental factors cause variation in the phenotypes of interacting species, the fitness outcome of their interaction, and reciprocal selection^5^. Consequently, coevolution can be absent/present (representing coevolutionary cold- and hotspots) and vary in form and direction on a continuous scale between mutualistic and antagonistic^2,3^. Different forms and directions of coevolution expose populations of coevolving organisms to different evolutionary trajectories: Mutualistic species interactions (both interactors benefit) drive mutualistic coevolution and related traits, such as plant-advertisement and pollinator-foraging traits, while antagonistic species interactions (one exploiter, one prey/host) drive antagonistic coevolution and related traits, such as plant defense (including the hypersensitive response to kill herbivore eggs^6^) and counter strategies of herbivores^2,7,8^. Despite decades of research, exactly how abiotic and biotic factors influence the type and direction of coevolution (*e*.*g*. mutualistic and antagonistic coevolution) is still poorly understood. Such knowledge is essential to understand if coevolving organisms can survive and co-adapt to rapid environmental change.

Our understanding is hampered, however, by a lack of experimental assessment of how abiotic and biotic factors interact with each other to produce the geographic mosaic of coevolution. Field studies on plant-insect coevolution show that climate^9,10^ and the presence of biotic agents^11–13^ display extensive spatial-temporal variation^14,15^ and are potentially important drivers of coevolution^16^. However, field studies struggle with identifying coevolution and the exact causal agents of its variation. Populations may simply respond similarly to environmental factors but without coevolving^17–20^, and differences between coevolving populations may be explained by temporal variation rather than variation in environmental factors^16^.

Experimental coevolution is a promising approach to investigate the contribution of specific environmental factors to the trajectories and diversification of coevolutionary outcomes^1,21^. Although previous studies on microbes provided important insights by showing that individual abiotic and biotic factors can influence coevolution, such insights cannot directly be applied to multicellular organisms because of their complex phenotypes with different constraints/trade-offs and tangled interactions. Unfortunately, experimental coevolution has so far not been utilized in plant-insect systems^11^. Still, experimental evolution has proven effective in showing strong interactive effects of abiotic and biotic factors on plant evolution. For example, soil composition and biotic agents (pollinators, herbivores) interactively impacted divergence in floral- and resistance traits of *Brassica rapa* plants^22–24^. Therefore, experimental coevolution with plants and insects is a promising avenue for testing existing hypotheses and providing new testable hypotheses about how abiotic and biotic factors collectively influence divergent coevolution.

We investigated if a combination of biotic and abiotic factors together could rapidly create a mosaic of coevolution (hypothesis 1), and thereby experimentally test a major prediction of the GMTC. We tested whether temperature and the presence of mutualistic bumblebee pollinators influenced experimental coevolution between a plant and its mixed mutualistic-antagonistic butterfly pollinator. The butterfly *Pieris rapae* is a generalized flower visitor and acts as efficient pollinator of *Brassica rapa* plants but also oviposits on plants from which the Brassicaceae-specialized caterpillars feed^5,25^.

According to the GMTC, the mutualistic-antagonistic outcome of coevolution depends on biotic factors such as the composition of the interacting community ^3,12,26^. We expect that the presence of bumblebees reduces the mutualistic value of butterflies for plants: even though both are efficient pollinators of *B. rapa*, bumblebees do not inflict costs of herbivory. Hence, the presence of bumblebees is expected to shift plant-butterfly coevolution to be more antagonistic (hypothesis 2)^12^.

The mutualistic-antagonistic outcome of coevolution can also depend on abiotic factors such as temperature ^9,27,28^. Elevated temperatures can reduce the mutualistic value of butterflies by reducing pollination efficiency and plant tolerance to herbivory, reducing the fitness benefit of butterfly visitation. Hence, elevated temperature is expected to shift plant-butterfly coevolution to be more antagonistic (hypothesis 3)^5^.

Combinations of environmental factors can change the mutualistic-antagonistic outcome of coevolution differently compared to individual factors, but specific outcomes are hard to predict^29,30^. Hence, we expect elevated temperature to alter the antagonistic coevolution triggered by bumblebees (hypothesis 4), but the precise direction is unclear. Potentially, the combination of bumblebee presence and elevated temperatures will reduce the shift towards more antagonistic plant-butterfly coevolution because multiple environmental factors can reduce antagonistic coevolution due to trade-off dynamics^27,28^. Exploring this question will lead to new testable hypotheses about how global warming will alter the geographic mosaic of coevolution.

To experimentally test these hypotheses, we performed experimental coevolution for 6 generations. Our controlled greenhouse environment allowed us to manipulate realistic ecological abiotic and biotic factors in a full-factorial manner, which is essential for detecting causality of individual factors and their interactions^31^. During the experiment, plants and butterflies were kept in cages (2.5×1.8×1.2m) in greenhouse cabins with different temperatures: “ambient” (23±2°C) and “hot” (27±3°C with 24h of 30±3°C per week); and in different biotic/pollination environments: no pollinators with plants being hand-pollinated (treatment referred to as: H), presence of coevolving butterflies (treatment referred to as: Coevo), presence of coevolving butterflies and non-coevolving bumblebees (treatment referred to as: CoB). The temperatures approximate current and predicted future mean temperatures in late spring/early summer in northern Switzerland (CH2018 model^32^). The hand pollination treatment was a greenhouse control to see whether the temperature environments could cause phenotypic divergence through differential plant mortality, fertility, and germination. All biotic treatments were replicated in both temperature environments, and all biotic-temperature combinations were replicated twice with 49 plants and 20-25 butterflies each (except H). Plant seed production resulting from insect pollination and number of eggs oviposited by butterflies contributed proportionally to each new generation of plants and butterflies. After 6 generations, we used generation 7 to reduce maternal effects: all plants were grown under ambient temperatures and hand-pollinated, while butterflies were exposed to stock rearing conditions. For the 8^th^ and final generation, we grew 36 plants from randomly selected mother plants of generation 7 from each treatment- and replicate combination together with 18 individuals of their respective butterfly lines or stock rearing (henceforth: control) butterflies. Afterwards, we measured plant and butterfly phenotypes, flower visitation and oviposition, and fitness in the ambient environment.

## Results

### Coevolution control

To separate coevolutionary and single-sided evolutionary changes, we included a one-sided-evolution control in the ambient environment where plants were exposed to non-evolving control butterflies in each generation^1^. Plants evolved in response to selection by butterflies, but butterflies did not, preventing coevolution. Indeed, we found significant differences in multivariate and individual trait divergence between plants evolving with non-evolving control butterflies and plants coevolving with butterflies (Coevo-plants) (supplementary results, Fig. S1, Table S1-S2). We also show that Coevo-butterflies significantly diverged from control butterflies (see below) suggesting butterflies evolved in response to coevolving plants rather than to greenhouse conditions. Together with our previous work showing reciprocal selection between plants and butterflies^5^, this collectively suggests plants and butterflies were indeed coevolving in our coevolution treatments.

### Hypothesis 1: Effect of combined abiotic and biotic factors on phenotypic divergence

We expected that the combination of abiotic and biotic factors would cause different co-evolutionary trajectories than each factor alone or the sum of both. We tested this by comparing multivariate phenotypic differences using linear discriminant analysis (LDA) combining all phenotypic traits for coevolving plants and butterflies. We found unique coevolutionary trajectories for plants and butterflies depending on the combination of abiotic and biotic factors which proves a major prediction of the GMTC. Multivariate phenotypic differences were most explained by the combination of temperature and biotic treatment, and less so by their individual effects (Table S1). For plants, the LDA showed separation of hand-pollinated control plants (H-plants) in both environments from all other plants (Fig. 1A). Plants coevolving with butterflies under ambient temperatures (ambient-evolved Coevo-plants) and plants coevolving with butterflies and bumblebees under elevated temperatures (hot-evolved CoB-plants) clustered together, and the same was true for ambient-evolved CoB-plants and hot-evolved Coevo-plants. For butterflies, the LDA showed hot-evolved CoB-butterflies far from control and Coevo-butterflies of both environments, while ambient-evolved CoB-butterflies differed the least from control butterflies (Fig. 1B).

**Figure 1.**
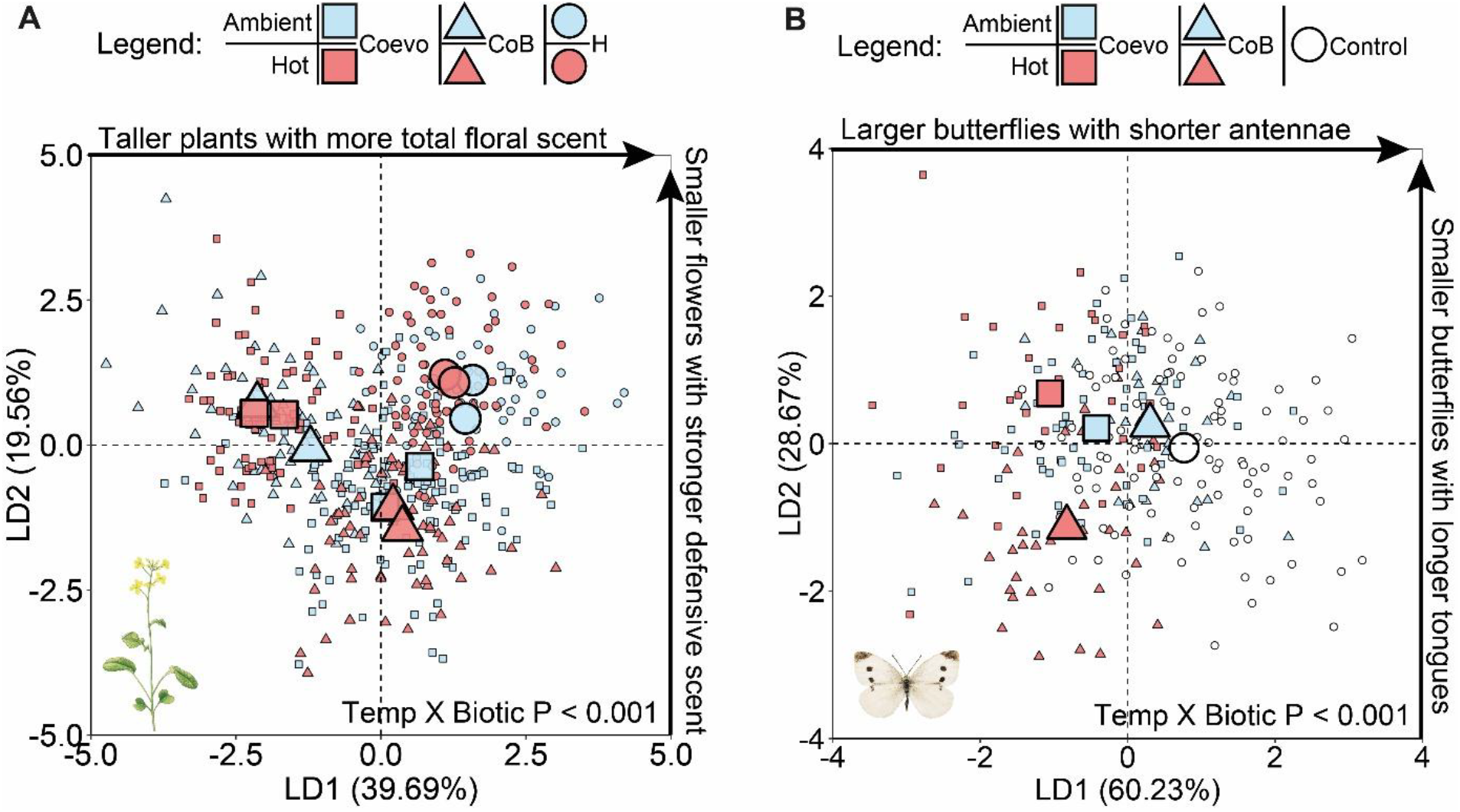
Phenotypic divergence of plants (A) and butterflies (B) after 6 generations of selection under different temperature and biotic/pollination environments. Blue = ambient temperature, red = hot temperature, H = hand pollination (circles in A), Control = *Pieris*-control (circles in B), Coevo = coevolution *Pieris*-only (squares), CoB = coevolution *Pieris* + bumblebees (triangles). Linear discriminant analyses included all plant phenotypic traits (n=19) and all butterfly phenotypic traits. For plants, double centroids per treatment combination represents replicates A and B. For butterflies, only one replicate was used due to replicate extinctions except for ambient Co, for which replicates were grouped to keep treatments comparable. P-values are based on permutational multivariate analysis of variance. Each temperature-biotic environment combination consisted of 68-144 plants and 36-100 butterflies. Characters with high loadings for LD1 and LD2 are shown along each axis.

### Hypothesis 2: Bumblebee presence led to more antagonistic coevolution

To assess if community composition changes the outcome of mutualistic-antagonistic coevolution, we tested if the presence of bumblebees led to more antagonistic coevolution (hypothesis 2). Overall, we found both mutualistic and antagonistic coevolution when plants and butterflies coevolved without other pollinators (Fig. 2): the evolution of pollinator-attraction traits (increased trait values of 5 traits compared to H-plants) support that plant-butterfly coevolution led to mutualistic coevolution, while reduced plant visitation, increased herbivore resistance, the evolution of (few) plant-resistance traits (increased trait values of 5 traits compared to H-plants) and smaller butterflies suggest simultaneous antagonistic coevolution (supplementary results, Table S3-S6). In the presence of bumblebees, we found a strong increase in plant herbivore resistance (plants received fewer eggs, induced more hypersensitivity response, suffered less fruit damage and caterpillar performance was strongly reduced), plants evolved traits related to defense (increased trait values of 5 traits compared to H- and Coevo-plants), and butterflies had reduced fitness, *i*.*e*. laid fewer eggs (Fig. 3, supplementary results, Table S3-S6). Together, these findings support hypothesis 2.

**Figure 2.**
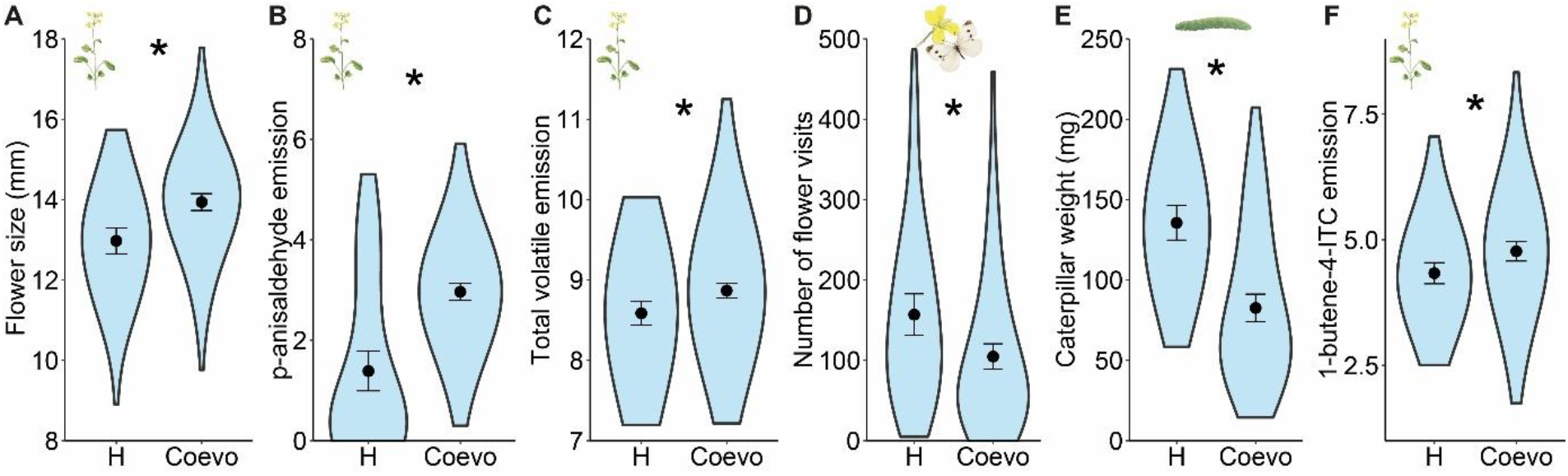
Plant-butterfly coevolution led to mutualistic and antagonistic coevolution. Plant-butterfly interactions and plant traits associated with mutualistic and antagonistic coevolution after 6 generations of selection with or without coevolving butterflies. Evolved traits associated with mutualistic coevolution were flower size (**A**), the emission of p-anisaldehyde (**B**), and total volatile emission (**C**). Evolved interactions and traits associated with antagonistic coevolution were number of flower visits received by plants (**D**), caterpillar weight (**E**), and the emission of 1-butene-4-ITC (**F**, ITC = isothiocyanate). All floral scent emission was log-transformed and in pg/f/l/h. H = hand pollination, Coevo = coevolution *Pieris-*only. Dots represent means, error bars are the 95% confidence interval of the means, colored areas show smoothened density traces: the distribution of data points where width represents data point density. Asterisks indicate significant differences at P ≤ 0.05. Each treatment consisted of 68-144 plants.

**Figure 3.**
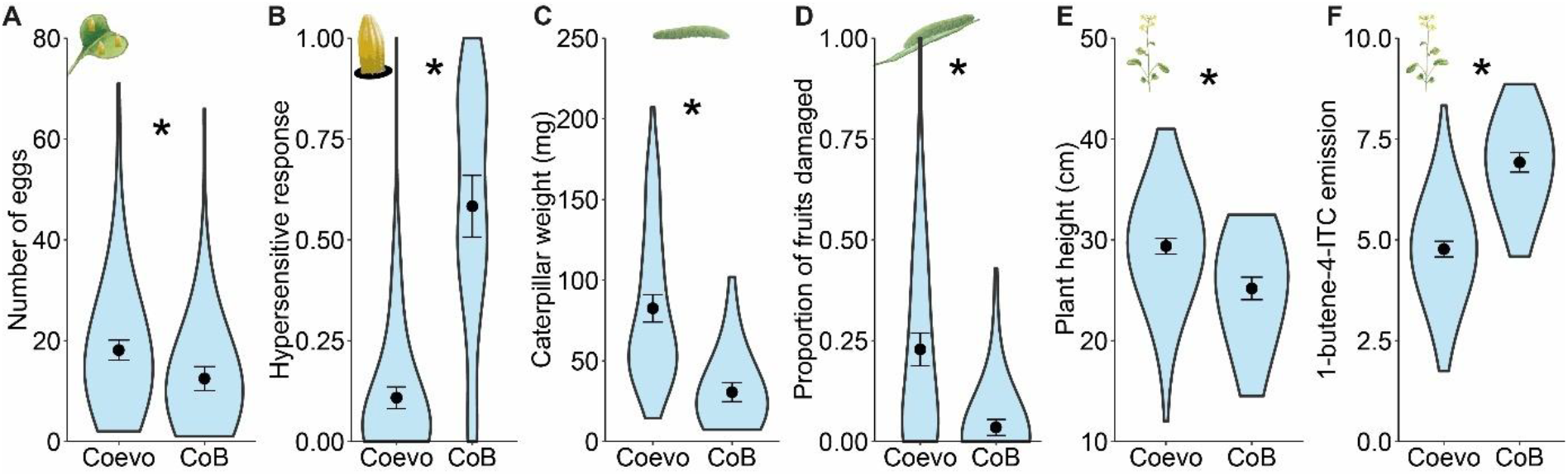
The presence of bumblebees led to more antagonistic coevolution. Plant-butterfly interactions and plant traits associated with antagonistic coevolution after 6 generations of selection with coevolving butterflies and with or without bumblebees present. Evolved interactions and traits associated with antagonistic coevolution were plant resistance: oviposition (**A**), hypersensitive-response as proportion of the number of eggs displaying hypersensitive response (eggs surrounded by necrotic plant tissue) divided by the total number of eggs (**B**), and caterpillar weight (**C**); proportion of fruits damaged as the number of fruits damaged divided by the total number of fruits (**D**), plant height (**E**), and the emission of 1-butene-4-isothiocyanate (**F**; emission log-transformed and in pg/f/l/h, ITC = isothiocyanate). Coevo = coevolution *Pieris-*only, CoB = coevolution *Pieris* + bumblebees. Dots represent means, error bars are the 95% confidence interval of the means, colored areas show smoothened density traces: the distribution of data points where width represents data point density. Asterisks indicate significant differences at P ≤ 0.05. Each treatment consisted of 68-144 plants.

### Hypothesis 3: Elevated temperature led to more antagonistic coevolution

To assess if abiotic factors can drive novel coevolutionary trajectories, we tested if elevated temperature led to more antagonistic coevolution (hypothesis 3). When plants and butterflies coevolved alone under elevated temperature, we found increased plant resistance (plants received fewer eggs, induced more hypersensitive response, and caterpillars performed worse) and plants evolved traits related to defense (increased trait values of 6-8 traits compared to H- and ambient-evolved plants, Fig. 4A-D, supplementary results, Table S7-S8). Plants experienced reduced flower visitation efficiency (lower increase in seed production per butterfly visit) and negative effects of herbivory on seed production (Fig. 4E, S3), likely causing reduced plant reproduction (supplementary results, Fig. S2, Table S10). Butterfly fitness was reduced, and butterflies became smaller (supplementary results, Table S9). Together, these findings support hypothesis 3 that elevated temperature led to more antagonistic plant-butterfly coevolution. However, an increase in flower visitation and the evolution of plant pollinator-attraction (increased trait values of 4-5 traits compared to H- and ambient-evolved plants) and butterfly-flower-foraging traits (increased trait values of 2 traits compared to H- and ambient-evolved plants) also indicates mutualistic coevolution (supplementary results, Table S7-S9).

**Figure 4.**
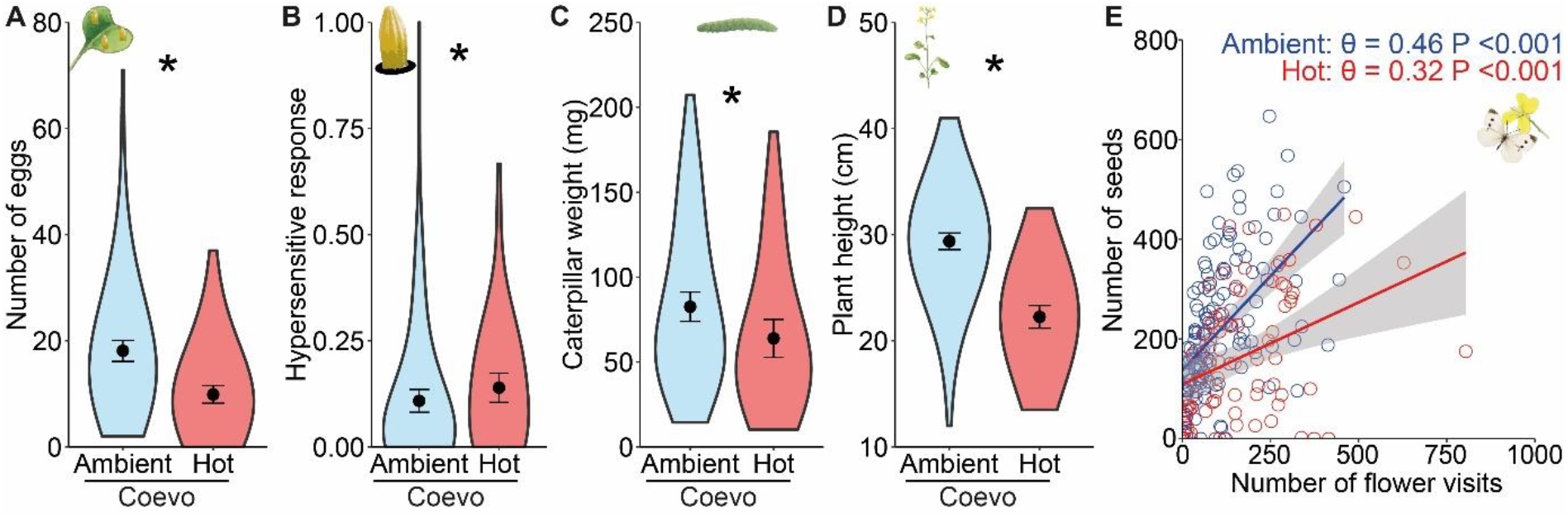
Elevated temperature led to more antagonistic coevolution. Plant-butterfly interactions and plant traits associated with antagonistic coevolution after 6 generations of selection with or without coevolving butterflies under ambient or elevated temperature. Evolved interactions and traits associated with antagonistic coevolution were plant resistance: oviposition (**A**), hypersensitive-response as proportion of eggs that show necrotic tissue (**B**), caterpillar weight (**C**); plant height (**D**), and reduced pollination efficiency as the relationship between seed production and number of flower visits (**E**). Blue = ambient temperature, red = hot temperature, H = hand pollination, Coevo = coevolution *Pieris-*only. Dots represent means, error bars are the 95% confidence interval of the means, colored areas show smoothened density traces: the distribution of data points where width represents data point density. Asterisks indicate significant differences at P ≤ 0.05, ns = non-significant. Each treatment consisted of 68-144 plants.

### Hypothesis 4: Combined bumblebee presence and elevated temperature changed coevolution

To assess if multiple environmental factors changed the mutualistic-antagonistic outcome of coevolution differently compared to individual factors, we tested if the combination of bumblebee presence and elevated temperature altered antagonistic plant-butterfly coevolution (hypothesis 4). When plants and butterflies coevolved under elevated temperatures in the presence of bumblebees, we found lower plant resistance (plants received more eggs, induced less HR, and caterpillars performed better, leading to more fruit damage), reduced evolution of plant defense traits (decreased trait values of 2-4 traits compared to H- and ambient-evolved plants), and tolerance to herbivory did not evolve (Fig. 5A-D, supplementary results, Fig. S3, Table S8,S10). Plant reproduction was reduced, while butterfly fitness was not (supplementary results, Fig. S2, Table S9-S10). These findings suggest that the combination of bumblebee presence and elevated temperatures led to less antagonistic coevolution. Simultaneously, few pollinator-attraction and butterfly-foraging traits increased (increased trait values of 2-3 traits compared to H- and ambient-evolved plants) while other plant-attraction- and butterfly-foraging traits were reduced (decrease in trait values of 3-4 plant traits compared to H- and ambient-evolved plants and 2-3 butterfly traits compared to control and ambient-evolved butterflies, supplementary results, Table S7-S9). Together with reduced flower visitation these changes indicate weak mutualistic coevolution. Overall, we provide support for hypothesis 4 that the combination of multiple environmental factors changes mutualistic-antagonistic coevolution differently compared to individual factors (Table S11-S14).

**Figure 5.**
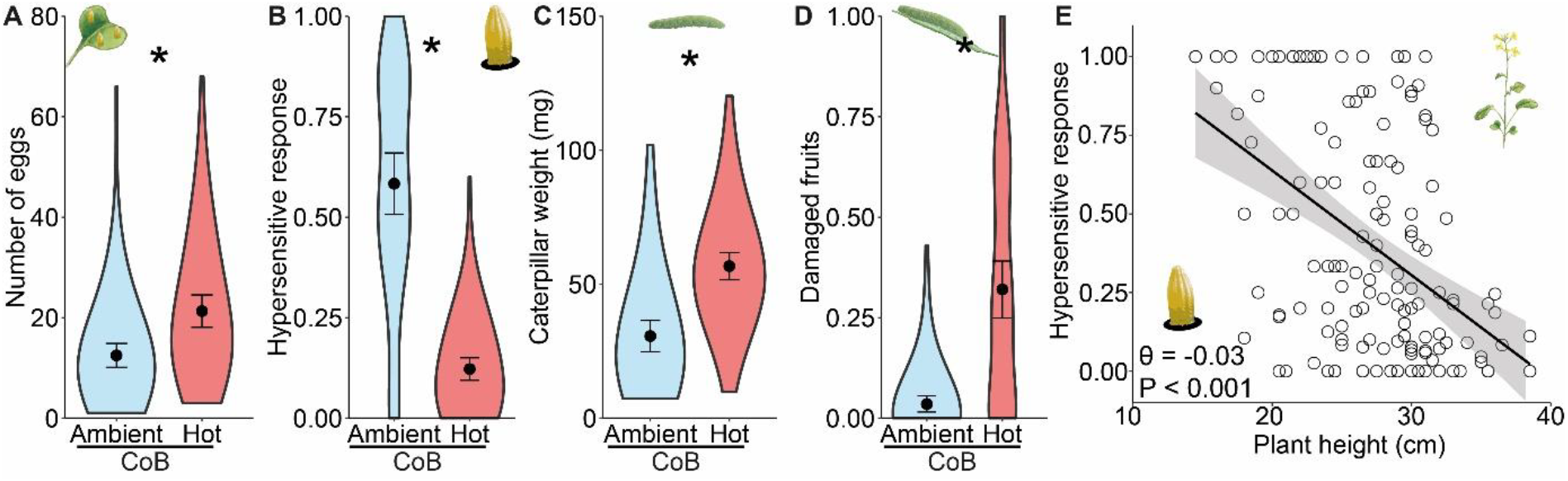
The combination of bumblebee presence and elevated temperature led to less antagonistic coevolution. Plant-butterfly interactions and butterfly traits associated with less antagonistic/more mutualistic coevolution after 6 generations of selection with or without bumblebees present and under ambient or elevated temperature. Evolved interactions and traits associated with less antagonistic coevolution were oviposition (**A**), hypersensitive-response as proportion of eggs that show necrotic tissue (**B**), caterpillar weight (**C**), and fruit damage as number of damaged fruits divided by total number of fruits (**D**). Negative correlation between hypersensitive response and plant height (**E**) for CoB-plants (ambient-a nd hot-evolved in black) suggests costs of HR. Blue = ambient temperature, red = hot temperature, CoB = coevolution *Pieris* + bumblebees. Dots represent means, error bars are the 95% confidence interval of the means, colored areas show smoothened density traces: the distribution of data points where width represents data point density. Asterisks indicate significant differences at P ≤ 0.05. Each treatment consisted of 68-144 plants.

## Discussion

To provide evidence for the GMTC, one needs to show that geographic structure is important to understand the dynamics of coevolution^33^. Geographic structure in coevolution is generated by geographic variation in environmental conditions leading to variation in phenotypes, fitness interactions and reciprocal selection between interacting species^3,5^. So far, evidence for the GMTC has mostly come from studies on geographic variation in trait correlations among interacting species^3,11,20^ and few experimental studies on microbes^27,28,34,35^. Without experimental validation however, trait correlations are neither necessary nor sufficient to show coevolution^17–20^. Such validation is a daunting task that requires decades of research and has only been performed in a few systems^36–38^. In nature, temperature and community context vary geographically^14,15^ and hence our mosaic imitates ecologically relevant conditions. Thereby, we show “geographic” variation in plant-butterfly coevolution and thus provide experimental proof for the overall GMTC. Our findings are relevant for many coevolving organisms because temperature and community context are common environmental factors that influence the evolutionary dynamics of a variety of interactions (competition, predation, symbiosis) engaged in by microbes, plants, insects and vertebrates^3,27,28,34–36,38^.

A major hypothesis of the GMTC is that novel coevolutionary dynamics emerge when environmental factors induce local variation in the fitness outcome of interspecific interactions and move the interaction along the continuous scale between mutualistic and antagonistic^3,39^. Primary support for this hypothesis came from seminal work on the *Lithophragma-Greya* system. *Greya* moths pollinate flowers of *Lithophragma* plants but also lay eggs in the flowers from which caterpillars hatch that later feed on the seeds, very similar to the *Brassica*-*Pieris* system we use here. When only *Greya* moths are present in the environment, these moths act as mutualists of *Lithophragma* plants by providing pollination despite their seed predation^26,37,40^. However, when other pollinators are present (flies and bees) the value of *Greya* moth pollination decreases, and the interaction becomes antagonistic due to the seed predation. Interestingly, *Lithophragma* plants abort more seed capsules with *Greya* eggs when other pollinators are present^12^, which is comparable with *Brassica* plants killing more *Pieris* eggs with the hypersensitive response (HR) in the presence of other pollinators that we observed in our experiment. This confirms the causality of community context impacting coevolutionary dynamics. Community context was hypothesized to drive geographic variation in coevolution because the presence of other pollinators varies geographically and can affect the fitness outcome of the plant-moth interaction^12,26,37,40^. Although this hypothesis gained empirical support from field studies in various systems over the years^11,13,36,38^, it was never experimentally proven, except in few microbial studies^35,41^. We hereby provide experimental proof that community context can drive novel coevolutionary trajectories.

Abiotic factors can potentially drive coevolutionary dynamics, but this has received little attention^42^. Field studies on camellia trees and seed-feeding weevils, and flowering shrubs and hawkmoth pollinators show latitudinal variation in (co)evolved traits and fitness, suggesting that climatic factors drive variation in coevolution^9,10,43^. Although temperature has been proposed as one of the driving factors^9,43^, climatic variables are almost impossible to separate for each other in the field and need experimental validation. In previous work, we showed experimentally that temperature changes reciprocal selection between our plants and butterflies^5^. Our current study extends on that work showing that temperature leads to phenotypic co-divergence and antagonistic coevolution and thereby provides experimental evidence that abiotic factors can drive novel evolutionary trajectories. Investigating the effects of abiotic factors on coevolution is especially relevant given the rapid changing climate coevolving organisms are exposed to globally.

Single environmental factors affect coevolving species directly as well as species responses to other environmental factors leading to yet unpredictable combined effects^29,30^. These unpredictable effects of combinations of abiotic and biotic factors are likely a general component of most coevolving systems but have so far not been shown experimentally^36^. While one field study showed that the combination of community context and climatic variation best predicted geographic variation in toxicity-resistance coevolution between garter snakes and newts, interactive effects were not separated from individual effects^38^. Here, we show experimentally that the combined effects of temperature and co-pollinator presence on plant-butterfly co-divergence differ from their individual effects. Interestingly, antagonistic plant-butterfly coevolution seemed less strong in environments with combined compared to individual factors. When bumblebees are present, heat stress may reduce antagonistic coevolution due to trade-off dynamics or new/reduced selection pressures^5,27,28^. For example, elevated temperatures reduced the evolution of HR when bumblebees were present likely due to resource trade-offs: HR seems costly for plants^44,45^, as suggested by a negative correlation between growth (height) and HR in our plants (Fig. 5E). Thus, individual factor-driven shifts in mutualistic-antagonistic coevolution can be negated by the presence of other environmental factors; a shift towards more antagonistic plant-butterfly coevolution due to the presence of bumblebees was reduced by heat stress and vice versa.

Our controlled greenhouse environment allowed us to 1) manipulate the presence of coevolution, which is essential to separate coevolutionary and single-sided evolutionary changes, 2) manipulate abiotic and biotic factors in a full-factorial manner, which is essential for detecting causality of individual factors and their interactions^31^. Simultaneously, our greenhouse environment did not reflect the complexity of environmental factors coevolving plants and butterflies are exposed to in nature, many of which likely influence coevolution. In addition, the GMTC hypothesizes three processes to be the primary drivers of coevolutionary dynamics: hot and cold spots, selection mosaics, and trait remixing^20^. Our study did not include trait remixing via gene flow and metapopulation dynamics. Since trait remixing can constrain or promote divergent coevolution^20^, it is hard to predict how trait remixing by for example gene flow would influence our result. Nevertheless, our study likely reflects ecologically-realistic outcomes: our system is relatively similar to the *Greya*-*Lithophragma* system and we observe similar ecological (and evolutionary) outcomes as identified in the field: increased antagonistic coevolution in the presence of co-pollinators including the increased disposal of herbivore eggs^12,26,37,40^.

Global warming will have profound effects on the geographic mosaic of coevolution. In addition to rising temperatures, global warming causes range shifts in most organisms and facilitates species invasions and extinction events^30,46^. Hence, coevolving organisms will experience changes in climate, biotic communities, and their combination throughout their geographic ranges. Our study shows that such changes in abiotic and biotic factors can lead to substantial changes in the form and direction of coevolution, leading to unique coevolutionary trajectories^3^. We show that global warming can increase the rate of antagonistic coevolution by changing temperature or introducing novel biotic agents, while reducing rates of antagonistic coevolution by simultaneously changing these factors^3,27,28^. Reduced rates of antagonistic coevolution can have consequences for the persistence of coevolving populations^47^. For example, reduced antagonistic plant-herbivore coevolution may lead to reduced adaptation of plants to (specialized) herbivores. Without plant adaptation, (specialized) herbivores may reduce plant fitness and consequently population sizes, leading to a reduction in genetic variation, adaptive potential and eventual extinction^47,48^. Indeed, experimental coevolution with competing bacteria showed that heat stress led to reduced evolution of the competitive ability of one of the bacteria, *i*.*e*. reduced antagonistic coevolution, ultimately leading to its extinction^49^. Investigating effects of global warming on coevolution in a community context will increase our understanding of how global warming will impact local co-adaptation, evolutionary rescue and potential extinctions^50^.

Overall, we confirm experimentally that biotic (community composition) and abiotic (temperature) factors can drive novel coevolutionary dynamics. We show that experimental coevolution is an efficient tool to verify the importance of environmental factors on coevolution indicated by field studies, while also providing novel targets (the combined effects of temperature and community context) for field evaluation. Ultimately, combining experimental and field studies will help understand how current rapid environmental change will influence one of the major driving forces of earth’s biodiversity, the geographic mosaic of coevolution^3,36^.

## Materials and Methods

### Model system

We used fast-cycling *Brassica rapa* L. plants for our experimental coevolution. Fast-cycling plants (Wisconsin Fast Plant, Caroline Biological Supply, Burlington) are annual and mostly self-incompatible. They are ideal for experimental coevolution because of their high genetic diversity, short generation time, and interactions with antagonistic and mutualistic organisms including the pollinating herbivore *Pieris rapae* L. and the bumblebee pollinator *Bombus terrestris* L.^22–24,51,52^. Pollinator-attraction traits and plant-defense traits are relatively well understood in this system (see supplementary information).

To ensure high genetic diversity in our butterfly rearing, we collected butterflies and eggs from 5 collection locations in Switzerland (Binz, Dübendorf, Einsiedeln, Sonnental, Zürich) and one in the Netherlands (Renkum) in 2020 (see supplementary information). Bumblebee hives were purchased from Andermatt Biocontrol (Hummelvolk Bombus Maxi, Andermatt Biocontrol, Grossdietwil, Switzerland); two hives were purchased per generation and for each bioassay to control for hive-specific effects. See supplementary information for details on insect rearing.

### Experimental coevolution: the plant

For each of the 7 treatments, we used 98 full-sib plant families (1 plant per family, see supplementary information) divided into two replicates (A and B) of 49 families as starting populations. Replicates A and B were subjected to the same treatments but never combined and thus evolved independently. This resulted in 14 populations of 49 plants. Eight days after sowing (see supplementary information for details on sowing) plants were moved to the greenhouses where biotic treatments were applied 24-27 days after sowing. For treatments with insects, plants were arranged in a seven-by-seven matrix inside a flight cage (2.5×1.8×1.2m). For Evo- and Coevo-plants, 25 female butterflies were released for 2 days. For CoB-plants, 6 individual worker bumblebees were released sequentially. Each bee was recaptured after visiting 4 plants. Simultaneously, 20 female butterflies were released for 2 days. Butterflies eclosed from pupae from the rearing were mated 2 days before the assays, starved 1 day before, and did not have any contact with plants. We counted the number of eggs oviposited on each plant 1-3 days after removing the butterflies, ranked plants for their egg load, assigned herbivory treatments (between 0-4 caterpillars per plant), and after few weeks collected and stored pupae in a refrigerator (±10°C) until further rearing (see supplementary for a more detailed description). For H-plants, we collected pollen with a make-up brush from 3-5 flowers of every plant within the replicate. We then pollinated 6 flowers per plant with this brush.

One month after applying the biotic treatments plants were moved to a different greenhouse and watering was stopped. When fruits were dry, seeds were counted by collecting all seeds of a plant on a white plate, photographing it and using ImageJ to count them. Based on the number of seeds, the contribution of each plant to the next generation was calculated: 49/(population sum of seeds/individual seed number). The next generation was sown accordingly and this allowed for natural selection and evolutionary change over 6 generations^23,24^. For hand pollinated plants (H-plants), we applied (cruder) selection on reproduction/survival: Upon successful reproduction, a plant contributed one individual to the next generation. Plants with unsuccessful reproduction (mortality, fertility/germination problems, etc.) were replaced by another randomly selected plant which reproduced successfully. This was done because our method of hand pollination (6 flowers) would likely not result in selection on reproduction via resource-limiting mechanisms or fertility for plants producing >20 flowers (most plants) and hand pollinating more flowers was not feasible. After 6 generations of experimental coevolution, we used generation 7 to reduce maternal effects: all plants were grown in the ambient environment and hand pollinated as described above, except 10 flowers were pollinated instead of 6. The resulting seeds were used to grow the 8th and final generation and check for coevolutionary changes.

### Experimental coevolution: the butterfly

We set up 8 butterfly lines from the collected pupae of the treatments Coevo and CoB, replicate A and B, ambient and hot temperatures. Butterflies experience selection alongside the plants and should evolve over multiple generations just like the plants^22,23,53^. Because we had to cover >4 weeks between plant generations, we reared an additional “in-between” generation for the butterflies between two plant generations. Butterfly development can be slowed down in the refrigerator (±10°C) but no longer than 3/4 weeks (pers. obs. Q. Rusman). Rearing conditions were similar as stock rearing conditions (see supplementary information), and hence we expected no selection on the lines other than rearing/greenhouse related. During plant generation 7, no insects were used, and butterflies underwent additional in-between generations, until plant generation 8 was ready.

### Evolutionary divergence of coevolved population

To investigate if plant-butterfly coevolution was influenced by temperature and bumblebee presence, we grew 36 plants (generation 8) from randomly selected mother plants (generation 7) from each treatment together with their respective butterfly lines or control (rearing) butterflies. Because the hot temperature butterfly lines were doing poorly between plant generation 7 and 8, we merged replicate A and B to prevent extinction and butterflies from the merged line (hot Coevo- or CoB-butterflies) were used to interact with plants from replicate A and B of the matching treatments. Because ambient CoB-butterfly line A went extinct between plant generation 7 and 8, we used line B in combination with plants from replicate A. To synchronize plants and butterflies, treatments were assigned to one of four cohorts separated by two weeks. Sowing, potting, and keeping the plants was done similar as described above; we kept all plants in the ambient environment to focus on genetic phenotypic and behavioral changes and exclude temperature-induced plasticity.

To investigate if coevolution in different environments led to phenotypic divergence, we measured plant and butterfly phenotypic traits likely involved in coevolution^5^. For plants, we measured height, leaf and flower number, flower size, nectar, and flower scent 21-28 days after sowing. For butterflies, we measured weight, size (wing area), tongue length, antennal and club lengths after butterfly visitation assays (see below). See the supplementary information for detailed descriptions of trait measurements.

#### Plant-butterfly interactions

To investigate if (co)evolution in different environments led to different plant-*Pieris* interactions, we performed butterfly visitation assays. Assays were carried out for 2 days, 29-32 days after sowing. Plants were arranged in a six-by-six matrix inside a flight cage (2.5×1.8×1.2m). 18 female butterflies of the matching lines or the rearing were released in the flight cage for 2 days. Butterflies were mated 2 days before the assays, starved 1 day before, did not have any contact with plants, and were marked before release. Each day, butterflies were released between 10:00-12:00am and 13:00-16:00pm. We recorded every flower visitation and oviposition made by each butterfly. After the assays, butterflies were stored at −20°C for phenotyping. Egg counting and herbivory treatments were similar as described above. During egg and excess larvae removal, eggs were monitored for HR: hypersensitive response of the plant to kill the egg, visible as a ring of black necrotic plant tissues around the egg’s base^54^. HR was calculated by dividing the number of eggs displaying HR by the total number of eggs per plant. Caterpillars were weighed 16 days after oviposition.

#### Plant reproduction

To investigate if (co)evolution in different environments led to different plant reproduction, we measured fruit set, number, damage, and the number of seeds in total and per fruit. One month after butterfly pollination, plants were moved to a different greenhouse and watering was stopped. When fruits were dry, we counted, assessed damage, and collected them. Fruit damage was estimated by dividing the number of damaged fruits by the total number. Fruits were considered damaged when displaying any visible damage such as chewed edges, holes, and missing parts. After collecting, we opened the fruits, photographed the seeds on a white platter, and used the software ImageJ to count them. Fruit set was estimated by dividing the number of flowers by the number of fruits. Seeds per fruit were estimated by dividing the seed number by fruit number.

### Statistical analyses

We analyzed phenotypic divergence with permutational multivariate analysis of variance (PERMANOVA) and linear discriminant analysis (LDA) by combining all traits. Plant scent compounds were log transformed to approach normal distribution. For one-sided- and co-evolution, models contained evolutionary history and replicate as fixed factors. For interactive effects of temperature and biotic treatments, we added temperature, biotic treatment, their interaction, and replicate as fixed factors. We used Euclidian distances and 1000 permutations.

To compare effects of evolutionary environments on individual trait evolution and plant-*Pieris* interactions, we used (generalized) linear mixed models. For plants, we compared one-sided- and co-evolution using models containing evolutionary history (H, Evo, Coevo) as fixed factors and replicate as random factor. To test interactive effects of temperature and biotic treatments, models contained temperature, biotic treatment (H, Coevo, and CoB) and their interaction as fixed factors and replicate and plant identity (only for caterpillar weight) as random factors. For butterflies and to test for interactive effects of temperature and biotic treatments, models contained temperature, biotic treatment (control/rearing, Coevo, and CoB) and their interaction as fixed factors and replicate as random factor. For continuous variables we used models with a Gaussian distribution, in case the data did not follow a normal distribution we used a Gamma distribution with log link function (data without zeros) or a Tweedie distribution (data with zeros). For count variables, we used models with a Poisson or a negative binomial distribution to correct for overdispersion. For HR, we used an ordered beta regression model, and values ≥ 1 were set at 1.

To test effects of evolutionary environment on reproduction, we used (generalized) linear mixed models. All models contained temperature, biotic treatment, their interaction, number of butterfly visits, herbivory (number of caterpillars), and number of flowers (except for fruit set) as fixed factors and replicate as random factor. For number of fruits and seeds, we used models with a negative binomial distribution. For number of seeds per fruit, we used a model with a Gaussian distribution. For fruit set, we used an ordered beta regression model, and values ≥ 1 were set at 1. For damaged fruits, we used a model with a Tweedie distribution.

For all variables that tested interactive effects of temperature and biotic treatments, two sets of post hoc tests were performed: 1) comparison of biotic treatments within each temperature treatment, 2) comparison of temperature treatments within each biotic treatment. For butterflies, to compare biotic treatments in the hot environment we included “ambient rearing/control” as control. We carried out all statistical analyses in R (v4.1.3 × 64, 2022, The R Foundation for Statistical Computing Platform) and used packages are described in the supplementary information.

## Data availability

The raw data will be made available on figshare upon acceptance of the manuscript.

## Acknowledgements

We thank John N Thompson for comments on an earlier version of this manuscript, Corinne Hertäg for help during the floral scent analyses and quantification, Laura Dällenbach, Basil Züllig, Chiara Joy Caspers, and Lena Schneider for help with various experimental tasks, Rayko Jonas and Markus Meierhofer for taking care of the plants. This research was funded by the Swiss National Science Foundation (grant no. 310030_208158).

## Author contribution

Conceptualization: QR, TF, JT, FPS; Experimental work: QR, TF, JT; Data analyses: QR; Writing – original draft: QR; Writing – review & editing: QR, TF, JT, FPS.

