## Supplemental information for "Coevolution in a warming world: an experimental test of the geographic mosaic of coevolution"

### Materials and Methods

*Plant full-sib families*

To create plant families, we grew 300 plants from seeds supplied by Carolina Biological Supply Co. (Burlington, USA) and randomly assigned them in pairs. Five flowers from each plant were reciprocally hand-pollinated within these pairs, and the resulting seeds from both parents were mixed, forming one full-sib family.

*Butterfly stock rearing*

To ensure high genetic diversity in our butterfly rearing, we collected butterflies and eggs from 5 collection locations in Switzerland (Binz, Dübendorf, Einsiedeln, Sonnental, Zürich) and one in the Netherlands (Renkum) in 2020. We crossed every Swiss population (Binz, Dübendorf, Einsiedeln, Sonnental, Zürich) with the Dutch population (Renkum) by putting virgin females of each individual Swiss population together with virgin males of the Dutch population and vice versa. We then combined the two female-male hybrid combinations (Dutch female X Swiss male + Swiss female X Dutch male) of each individual Swiss and Dutch population combination, creating 5 hybrid lines (BinzXNL, DübXNL, EinXNL, SonXNL, ZürXNL). Subsequently, we did 3 rounds of random crossings between the hybrid lines resulting in 1 final population that contains genetic material from all 5 Swiss and the 1 Dutch population. Adult butterflies were kept in a rearing cage (90 x 60 x 60 cm) and provided with water, nectar, and flowering fast-cycling B. rapa plants to oviposit under greenhouse conditions (23 ± 1 ºC, 60% RH, 16h:8h, light:dark). Young larvae (L1-L3) were fed with flowering fast-cycling B. rapa plants and late-instar larvae (L4-L5) were transferred to savoy cabbage leaves (Brassica oleraceae var. sabauda).

*Bumblebee rearing*

Bumblebees were fed flowering fast-cycling *Brassica rapa* plants, and supplementary nectar (BioGluc, Biobest, Westerlo, Belgium) and pollen (Blütenpollen Multiflora, KoRo Handels GmbH, Berlin, Germany). To avoid pollen contamination during our experiment, plants were removed from the bumblebee cages two days prior to the bioassays. Supplementary pollen and nectar were removed 16 hs before bioassays.

*Plant sowing*

Every generation, plants were sown in multi-pot trays filled with a sowing mix consisting of equal parts sterilized sieved topsoil, compost (3 years-old), pumice (1-5 mm) and peat, and allowed to germinate in a phytotron (22±1ºC, 60%RH, 24h light). After 8 days, seedlings were re-potted in individual pots (7x7x8 cm) filled with standardized soil (Einheitserde, Germany) and moved to the greenhouses (60%RH, 16h:8h, light:dark). Because plants and butterflies developed faster in the hot environment, we divided treatments with coevolving butterflies into two cohorts (ambient and hot) which were sown two weeks apart and ambient treatments always first. For H- and Evo-plants, one of each replicate was randomly assigned to cohort 1 or 2.

*Plant herbivory treatments*

We counted the number of eggs oviposited on each plant 1-3 days after removing the butterflies. Plants were ranked for their egg load (rank 1 having the most eggs and rank 49 having the least eggs) and assigned to five herbivory treatments: Plants ranked 1-9 kept four caterpillars, 10-19 three caterpillars, 20-29 two caterpillars, 30-39 one caterpillar, and 40-49 zero caterpillars. We removed all excess larvae 7-9 days after oviposition. Larvae removal was done to simulate natural levels of egg and larvae survival, which can be as low as 0.1%[1], while ensuring herbivory proportional to oviposition and avoiding caterpillar amounts >4 which would kill the plant[2,3]. A few days later, we rechecked the caterpillar numbers per plant and fitted plastic reverse cones made from 3 transparent PVC A4-sized pages around each plant, including plants without caterpillars. These cones had an open top, with its border enveloped in cotton wool, preventing caterpillars from switching plants without hindering plant growth. We collected all pupae 3-5 weeks after oviposition and stored them in a refrigerator (±10°C) until further rearing. After the first pupae check, cones were removed from plants without caterpillars and for plants with caterpillars when all pupae were collected.

*Butterfly "in-between" generation rearing*

We set up 8 butterfly lines from the collected pupae of the treatments Coevo and CoB, replicate A and B, ambient and hot temperatures. Butterflies experience selection alongside the plants and should evolve over multiple generations just like the plants[4-6]. Because we had to cover >4 weeks between plant generations, we reared an additional "in-between" generation for the butterflies between two plant generations. Butterfly development can be slowed down in the refrigerator (±10°C) but no longer than 3/4 weeks (pers. obs. Q. Rusman). After pupae of each line (~100 per line, recovered from the experimental plants) were taken out of the refrigerator, they were put in rearing cages to eclose (similar conditions as described above and a separate cage for each line). Adults butterflies mated and were offered 2 flowering fast-cycling *B. rapa* plants for oviposition every 3-4 days (4 times, 8 plants in total). We standardized population sizes of the different lines during oviposition: If oviposition was high (>100 eggs in one day) plants were removed after 1 day, and if oviposition was low (<100 eggs in one day), plants were left for the full 3-4 days until the next batch of plants was presented. This yielded around 800 eggs in total for each line. All hatching caterpillars were reared until pupation under stock rearing conditions (see above). Each line generated 100-500 pupae each "in-between" generation. High mortality was mostly due to disease, which liquified caterpillars during development, and is a common problem in rearing *P. rapae* (pers. obs. Q. Rusman). All pupae were stored in the refrigerator until further use: 7-10 days before butterflies were needed for the experiment, all pupae were taken out of the refrigerator and allowed to eclose and mate under stock rearing conditions, from which 20 (CoB) - 25 (Coevo) mated female butterflies were selected and used in the subsequent butterfly visitation bioassays (see section below).

*Plant pollinator attraction and defense traits*

We considered the following traits pollinator attraction traits: (increased) flower number, (increased) plant height, flower size, nectar, and increased total floral scent emission and the

emission of p-anisaldehyde, phenylacetaldehyde, (E,E)-α-farnesene, (Z,Z)-α-farnesene. These traits often play a role in pollinator attraction[2,3,6,7].
We considered the following traits plant defense traits: (reduced) flower number, (reduced) plant height and (enhanced) emission of phenylethyl alcohol, 1-butene-4-isothiocyanate, methyl anthranilate, 3-hexenyl acetate, methyl salicylate, and indole. Reduced flower number, plant height and enhanced emission of phenylethyl alcohol, 1-butene-4-isothiocyanate, 3-hexenyl acetate, methyl salicylate, and indole can reduce (general) herbivore attraction[8-16], although 1-butene-4-isothiocyanate is attractive for *Brassica* specialist herbivores[9], including *P. rapae*[10]. Methyl anthranilate may act as plant defense by reducing egg hatching rate [11].

*Measuring plant phenotypic trait divergence: Non-scent flowering traits*
Between 21-24 days after sowing, we measured height and number of leaves for every plant. From three flowers per plant, we collected nectar and measured their diameter with a calliper. Nectar was collected with 1 µl capillaries (Blaubrand, Wertheim, Germany). To collect nectar, we gently pressed the capillary against the nectaries. We used the same capillary to extract nectar from the three flowers per plant. The measured amount was then divided by three to obtain the average nectar production per flower per plant. 28 days after sowing, we counted the number of open flowers. Nectar and flower diameter measures were averaged per plant.

*Measuring plant phenotypic trait divergence: Volatile collection and quantification*
Floral volatiles were collected and quantified via headspace sorption and by using a push-pull system [2,3]. Collections were done 21-25 days after sowing. We encased the main inflorescence of each plant in a glass cylinder with a silanized surface (sigmacote; Sigma-Aldrich, Buchs, Switzerland) on the inside. At the base, the glasses were sealed with two teflon plates, which only let the inflorescence stem through a hole (0.5 cm ⌀). Air was pushed through a side opening in the glass cylinder (fitted with an activated charcoal filter to purify the incoming air) and pulled through another opening, which contained a glass tube filled with an absorbent (30 mg of Tenax). Both pushing and pulling were performed by pumps at a rate of 110 ml/min. At the start of each collection, we counted the number of flowers encased within the glass cylinders. Collections lasted for 2 hours and were done between 11:00 - 13:00 and 14:00-16:00 h. One sample from an empty glass cylinder was collected as air control during each collection round. Quantification of volatiles was done by gas chromatography with mass selective detection (GC–MSD), see below. Because of potential contamination, we subtracted amounts collected of each compound in the air control from the samples per collection round. All volatiles were standardized in units of pg per flower per l sampled air per hour.

Samples were injected into a GC (Agilent 6890N; Agilent Technologies, Santa Clara, CA, USA) by a MultiPurpose Sampler (MPS, Gerstel, Mülheim, Germany) using a Gerstel thermal desorption unit (TDU, Gerstel) and a cold injection system (CIS; Gerstel). The GC was equipped with a HP-5 column (15 m length, 0.25mm ID, 0.25µm film thickness) and helium was used as a carrier gas at a flow rate of 2ml/min−1. Sampled volatiles were desorbed from the tenax by heating the TDU from 30 to 240°C at a rate of 60 °C min-1 and held at the final temperature for 5 min. Eluting compounds from the TDU were trapped in the CIS at -150°C. For injection, the CIS was heated to 250°C at a rate of 12°Cs-1, and the final temperature was held for 3 min. For compound identification and quantification, we used a mass selective detector (Agilent MSD 5975). Each compound was identified by comparing its mass spectra with those of the National Institute of

Standards and Technology (NIST) mass spectral library and synthetic standards of all compounds previously analysed on the GC-MSD system. Quantification (total ion counts) of the different compounds was done by using a calibration curve for target ions specific to the individual compounds.

*Measuring butterfly phenotypic trait divergence*

After defrosting, we weighed butterflies and then the head and wings were removed for measurements. The wing surface area was measured by gluing the front wings to a black sheet of paper using transparent sticky tape. We scanned the paper sheets with a printer (Canon imageRunner Advance DX) and the ensuing .jpg files were analyzed using the software ImageJ to calculate wing surface area (in cm2). We used a caliper to measure tongue, antennae and club lengths. Relative club length was calculated by dividing club length by antennae length.

*Statistical analyses*

For (G)LMMs, we used lme4[17], multcomp[18], lsmeans[19], and glmmTMB[20] packages, and we used F- and Wald chi-square tests to derive p-values. For PERMANOVAs, we used the vegan package[21]. For LDA visualization, we used the ggord package[22]. For violin plots, we used the ggplot2 package[23,24].

### Results

*Coevolution control*

Combining all plant phenotypic traits, we found significant divergence between plants from the hand pollination treatment (H-plants), one-sided evolution treatment (Evo-plants) and coevolution with butterflies but without bumblebees treatment (Coevo-plants) in the ambient environment (Table S1, Fig. S1). In the one-sided-evolution treatment more traits evolved and more strongly compared to the coevolution treatment: Compared to plants coevolving with butterflies (Coevo-plants), plants evolving with non-evolving control butterflies (Evo-plants) were larger (1.12-fold) with more flowers (1.33-fold) that had more nectar (1.56-fold) and emitted more of all scent compounds except hexenyl acetate (Table S2).

*Plant-butterfly coevolution led to mutualistic and antagonistic coevolution.*

To test if the presence of butterflies alone led to mutualistic and antagonistic coevolution, we compared plants and butterflies coevolving alone (Coevo-plants and butterflies) and hand pollinated plants (H-plants) and control butterflies. We found some evidence for mutualistic coevolution: Compared to H-plants, Coevo-plants evolved increased pollinator-attraction traits: plants had larger flowers (1.07-fold) with increased emission of six floral scent compounds including the known butterfly attractant p-anisaldehyde (1.74-fold) and the sum of all floral volatiles (1.31-fold; Fig. 2, Table S3). We also found evidence for antagonistic coevolution: Compared to H-plants, Coevo-plants received less flower visits (.91-fold), evolved increased herbivore resistance (*i.e.* caterpillars had reduced performance, .61-fold) and increased plant defense traits such as the emission of 1-butene-4-isothiocyanate (2.06-fold) and 3-hexenyl acetate (1.83-fold)(Fig. 2, Table S3-S4). Herbivory negatively affected reproduction for Coevo-plants but not for H-plants (Fig. S3, Table S6). Compared to control butterflies, Coevo-butterflies were lighter (.91-fold), had smaller wings (.91-fold) and antenna (.98-fold) (Table S5), potentially indicating antagonistic coevolution.

*The presence of bumblebees led to more antagonistic coevolution.*

To test if the presence of bumblebees led to more antagonistic coevolution, we compared plants and butterflies coevolving with the presence of bumblebees (CoB-plants and butterflies) and without bumblebees (Coevo-plants and butterflies). Compared to Coevo-plants, CoB-plants evolved increased herbivore resistance: plants received less eggs (.69-fold), induced more hypersensitivity response (HR; 6.22-fold), and caterpillar performance was strongly reduced (.37-fold) (Fig. 3, Table S4). In addition, CoB-plants had less fruit damage compared to Coevo-plants (.15-fold) (Fig. 3, Table S6). Contributing to the increased herbivore resistance are potential evolved traits related to resistance (reduced attraction): compared to Coevo-plants, CoB-plants were smaller (.86-fold) with smaller flowers (.96-fold) that emitted more scent compounds emission related to defense and less related to pollinator attraction (Fig. 3, Table S3). CoB-butterflies had lower reproduction compared to Coevo-butterflies (Fig. S2, Table S5), potentially indicating more antagonistic coevolution. This is not corroborated by changes in phenotypic traits however: CoB-butterflies were lighter (.94-fold) and smaller (.96-fold) than control butterflies, but heavier (1.08-fold) and larger (1.05-fold) than Coevo-butterflies, with longer antenna's (1.02-fold) (Table S5). These phenotypic changes suggest antagonistic coevolution for CoB-butterflies, but less severe than for Coevo-butterflies.

*Elevated temperature led to more antagonistic coevolution.*

To test if elevated temperature led to more antagonistic coevolution, we compared plants and butterflies that coevolved under elevated temperatures (hot-evolved Coevo-plants and butterflies) with hot-evolved control plants (H-plants) and butterflies, as well as hot- and ambient-evolved Coevo-plants and butterflies. Hot-evolved Coevo-plants evolved increased resistance compared to hot-evolved H-plants and ambient-evolved Coevo-plants: plants received less eggs (H-plants: 0.50-fold, ambient-evolved: .54-fold), induced more HR (ambient-evolved: 1.29-fold) and caterpillars performed worse (H-plants: 0.50-fold, ambient-evolved: .72-fold)(Fig. 4, Table S8). Hot-evolved Coevo-plants experienced reduced flower visitation efficiency (lower increase in seed production per butterfly visit) and negative effects of herbivory on seed production (Fig. 4, S3), potentially causing reduced reproduction: plants had lower fruit set, number of fruits, and number of seeds (Fig. S2, Table S10). Hot-evolved Coevo-plants evolved potential resistance traits: plants were smaller (H-plants: .76-fold, ambient-evolved: .76-fold) with less flowers (ambient-evolved: 0.80-fold) and increased scent emission related to defense (Table S7). Hot-evolved Coevo-butterflies laid less eggs (control butterflies: .53-fold, ambient-evolved: .52-fold), were lighter (control: .81-fold, ambient-evolved: .94-fold), and had smaller wings (control: .93-fold) (Table S9).

Contrary to hypothesis 3, hot-evolved Coevo-plants received more flower visits (H-plants: 1.49-fold, ambient-evolved: 1.72-fold) and butterflies evolved faster flower visitation speed (control butterflies: 1.37-fold, ambient-evolved: 1.59-fold)(Table S8-S9). Hot-evolved Coevo-plants evolved potential pollinator-attraction traits: plants had larger flowers (H-plants: 1.05-fold) with more nectar per flower (H-plants: 1.46-fold) and increased scent emission related to pollinator attraction (Table S6). Hot-evolved Coevo-butterflies evolved increased flower-foraging traits: longer clubs (control butterflies: 1.07-fold, ambient-evolved: 1.07-fold) (Table S9).

*The combination of bumblebee presence and elevated temperature led to less antagonistic coevolution.*

To test that the combination of bumblebee presence and elevated temperatures led to altered antagonistic plant-butterfly coevolution, we compared plants and butterflies that coevolved with bumblebees present (CoB-plants and butterflies) under elevated (hot-evolved) and ambient temperatures (ambient-evolved), and hot-evolved plants and butterflies with (CoB) and without (Coevo) bumblebees present.

We found reduced antagonistic coevolution: Hot-evolved CoB-plants evolved lower resistance: plants received more eggs (Coevo-plants: 2.15-fold, ambient-evolved: 1.71-fold), induced less HR (ambient-evolved: .18-fold) and caterpillars performed better (ambient-evolved: 1.85-fold) (Fig. 5, Table S8). Hot-evolved CoB-plants did not evolve tolerance to herbivory (ambient-evolved did, Fig. S3). Hot-evolved CoB-plants received more fruit damage (Fig. 5-S2), likely contributing to reduced reproduction: Hot-evolved CoB-plants had lower fruit set, number of fruits, number of seeds and less seeds per fruit compared to hot-evolved Coevo-plants and ambient-evolved CoB-plants (Fig. S2, Table S10). Hot-evolved CoB-butterflies reproduction was not reduced as compared to hot-evolved Coevo-butterflies and ambient CoB-butterflies (Table S9), suggesting less antagonistic coevolution.

We also found support for weak mutualistic coevolution to the combination of bumblebee presence and elevated temperatures compared to each factor individually: Hot-evolved CoB-plants received less flower visits (Coevo-plants: .45-fold, ambient-evolved: .62-fold), likely also contributing to the reduction in reproduction (Fig. S2, Table S8). Hot-evolved CoB-plants evolved reduced pollinator-attraction traits: plants evolved less nectar per flower (Coevo-plants: .61-fold, ambient-evolved: .63-fold) and reduced emission of total volatiles (Coevo-plants: .55-fold, ambient-evolved: .80-fold) and most scent compounds (Table S7). Hot-evolved CoB-butterflies evolved slower flower visitation speed (Coevo-butterflies: .47-fold, ambient-evolved: .65-fold) (Table S9). In contrast, hot-evolved CoB-plants evolved several pollinator-attraction traits: plants were larger (Coevo-plants: 1.31-fold, ambient-evolved: 1.26-fold) with more flowers (Coevo-plants: 1.53-fold, ambient-evolved: 1.26-fold) and higher emission of p-anisaldehyde (ambient-evolved: 1.80-fold) (Table S7). Hot-evolved CoB-butterflies evolved foraging traits: butterflies were heavier (Coevo-butterflies: 1.14-fold), had shorter tongues (ambient-evolved: .94-fold) and with longer absolute (ambient-evolved: 1.10-fold) and relative clubs (Coevo-butterflies: 1.05-fold, ambient-evolved: 1.12-fold) (Table S9). Shorter tongues and longer club length likely evolved because temperature-induced plasticity reduces floral scent emission and nectar amounts[2,3], and increased antennal clubs allow better detection of low floral scent[25,26], while reduced tongue length may be important to maintain foraging efficiency on open flowers with low nectar amount[27].

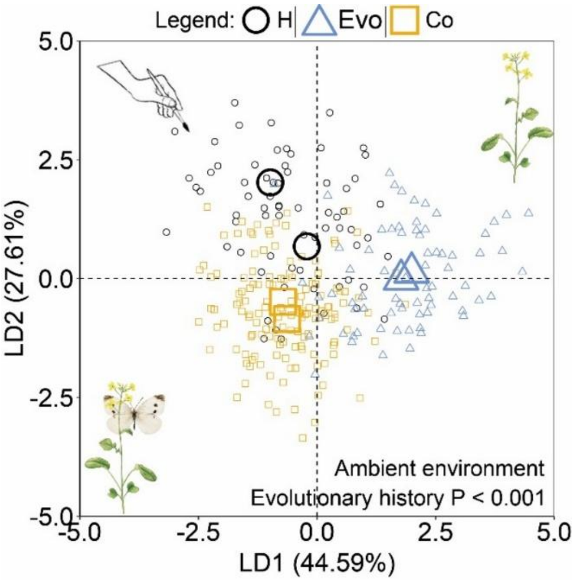


**Figure S1.** Phenotypic divergence of plants after 6 generations of selection under ambient temperature and different biotic/pollination environments. H = hand pollination, Evo = plant evolution only, Co = coevolution. Linear discriminant analyses included all plant phenotypic traits. Double centroids per treatment represent replicate A and B. P-values based on permutational multivariate analysis of variance. Each treatment consisted of 68-144 plants.

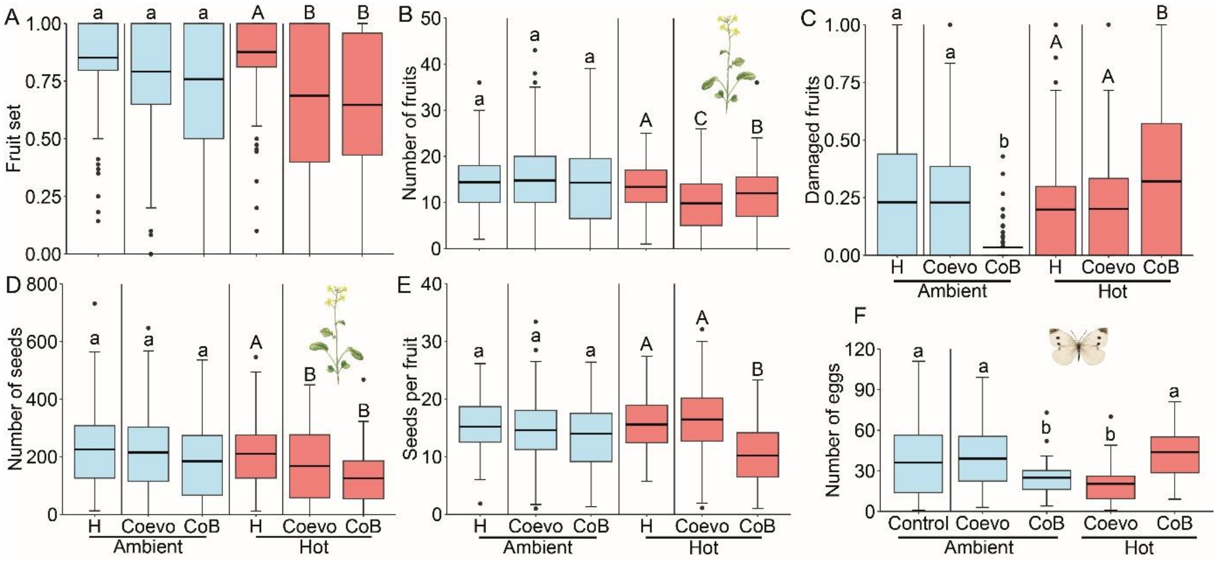


**Figure S2.** Reproduction of co-diverged plants (A-E) and butterflies (F) plants after 6 generations of selection under different temperature and biotic/pollination environments. Reproduction was measured in the focal environment: ambient temperature and butterflies as pollinators. Blue = ambient temperature, red = hot temperature, H = hand pollination (A-E), Coevo = coevolution *Pieris*-only, CoB = coevolution *Pieris* + bumblebees, Control = *Pieris*-control (F). Fruit set is the proportion of flowers that produced fruits (number of fruits divided by the number of flowers). Boxplots show mean (line). For plants, letters above bars indicate significant differences at P ≤ 0.05 based on Tukey's post hoc tests comparing biotic treatments within the ambient environment (small letters) and hot environment (capital letters). For butterflies, all treatments were compared. Each treatment consisted of 68-144 plants and 36-100 butterfly replicates.

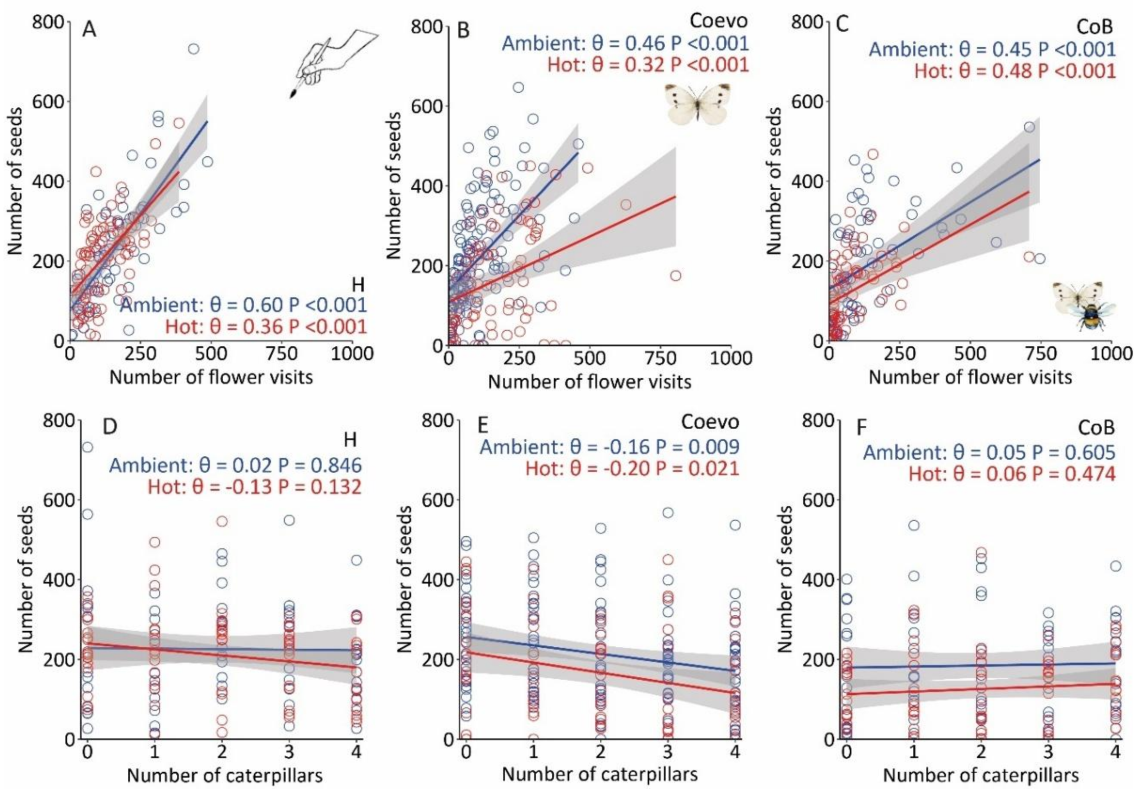


**Figure S3.** Relationship between seed production and number of flower visits (A-C) or number of caterpillars (D-F) of co-diverged plants and butterflies after 6 generations of selection under different temperature and biotic/pollination environments. Traits were measured in the focal environment: ambient temperature and butterflies as pollinators. Blue = ambient temperature, red = hot temperature, H = hand pollination, Coevo = coevolution *Pieris*-only, CoB = coevolution *Pieris* + bumblebees. Each treatment consisted of 68-144 plants.

**Table S1.** Output of permutational multivariate analyses of variance (PERMANOVA) showing effects of temperature and biotic environment on phenotypic divergence of plants and butterflies. All plant and butterfly phenotypic traits were included in the analyses. Bold values indicate results where P ≤ 0.05. Each treatment consisted of 68-144 plants and 36-100 butterflies.

| | Temperature | | | | Biotic | | | | T*B | | | | Replicate | | | |
|---|---|---|---|---|---|---|---|---|---|---|---|---|---|---|---|---|
| | df | F | *P* | $R^2$ | df | F | *P* | $R^2$ | df | F | *P* | $R^2$ | df | F | *P* | $R^2$ |
| Coevolution control | 2 | 50.99 | **<0.001** | 26.40 | - | - | - | - | - | - | - | - | 1 | 0.25 | 0.625 | 0.1 |
| Plant | 1 | 5.68 | **0.002** | 1.0 | 2 | 16.25 | **<0.001** | 5.5 | 2 | 25.40 | **<0.001** | 8.6 | 1 | 3.46 | **0.017** | 0.6 |
| Butterfly | 1 | 1.72 | 0.181 | 0.8 | 1 | 8.41 | **0.004** | 4.1 | 1 | 13.49 | **<0.001** | 6.5 | 1 | 5.70 | **0.012** | 2.8 |

**Table S2.** Trait values (mean ± SE) of evolved plant traits after 6 generations of selection under different biotic treatments in the ambient environment. H = hand pollination, Evo = one-sided evolution, Coevo = coevolution. Bold values indicate results where P ≤ 0.05 based on (generalized) linear mixed models (Biotic treatment). Letters indicate significant differences at P ≤ 0.05 based on Tukey's post hoc tests comparing biotic treatments. Scent emission is expressed as pg/flower/l/h. Each biotic environment consisted of 68-144 plants. ITC = isothiocyanate.

| | | | | Biotic treatment | | |
|---|---|---|---|---|---|---|
| | H | Evo | Coevo | $\chi^2$ | df | *P* |
| Flower number | 13.56±0.60a | 19.46±0.79b | 14.67±0.53a | 34.28 | 2 | **<0.001** |
| Leaf number | 5.90±0.18 | 5.97±0.18 | 5.61±0.14 | 1.36 | 2 | 0.506 |
| Plant height (cm) | 30.43±0.59a | 32.85±0.54b | 29.39±0.40a | 25.60 | 2 | **<0.001** |
| Flower diameter (mm) | 12.97±0.16a | 13.74±0.15b | 13.94±0.11b | 27.57 | 2 | **<0.001** |
| Nectar (µl) | 1.45±0.13a | 2.06±0.2b | 1.32±0.11a | 16.27 | 2 | **<0.001** |
| Total volatile emission | 6454.98±472.45a | 17572.02±1128.18c | 8440.33±607.64b | 124.1 | 2 | **<0.001** |
| Benzaldehyde | 3196.38±347.19a | 6074.56±719.3c | 4697.29±511.2b | 33.49 | 2 | **<0.001** |
| 1-butene-4-ITC | 117.69±18.83a | 421.34±61.09c | 242.81±37.92b | 56.20 | 2 | **<0.001** |
| Methyl benzoate | 144.67±15.69a | 445.17±56.75b | 117.01±8.53a | 131.54 | 2 | **<0.001** |
| Phenylethyl alcohol | 1.02±0.18a | 15.72±4.66b | 1.74±0.98a | 179.13 | 2 | **<0.001** |
| 2-Amino benzaldehyde | 1644.32±154.67a | 6036.23±513.03b | 1550.74±147.72a | 90.47 | 2 | **<0.001** |
| p-Anisaldehyde | 18.41±4.60a | 56.4±9.46c | 31.95±3.89b | 24.19 | 2 | **<0.001** |
| Methyl anthranilate | 150.38±17.06a | 579.48±62.73b | 131.26±11.72a | 120.26 | 2 | **<0.001** |
| (*Z*)-3-Hexenyl acetate | 99.55±10.6a | 134.35±16.01b | 182.58±32.26b | 20.29 | 2 | **<0.001** |
| Phenylacetaldehyde | 203.06±45.76a | 942.69±220.93b | 381.81±134.72a | 39.31 | 2 | **<0.001** |
| Benzyl nitrile | 148.91±22.55a | 513.37±46.59b | 173.16±14.83a | 64.00 | 2 | **<0.001** |
| Methyl salicylate | 135.69±15.26a | 322.26±28.23c | 204.49±29.31b | 33.59 | 2 | **<0.001** |
| Indole | 245.29±32.06a | 1297.47±103.81b | 341.4±29.28a | 114.22 | 2 | **<0.001** |
| (*Z,Z*)-α-Farnesene | 10.36±0.81a | 23.26±1.62c | 13.19±0.62b | 83.19 | 2 | **<0.001** |
| (*E,E*)-α-Farnesene | 339.23±36.68a | 709.71±52.4b | 370.9±21.28a | 61.76 | 2 | **<0.001** |

**Table S3.** Trait values (mean ± SE) of evolved plant traits after 6 generations of selection under different biotic treatments in the ambient environment. H = hand pollination, Coevo = coevolution, CoB = coevolution + bumblebees. A/D = attraction/defence, indication of which traits we consider attraction- and defense-traits based experience in the system and the literature (see supplementary methods above). Letters indicate significant differences at P ≤ 0.05 based on Tukey's post hoc tests comparing biotic treatments within the ambient environment. Scent emission is expressed as pg/flower/l/h. Each treatment consisted of 68-144 plants. ITC = isothiocyanate.

| | A/D | H | Coevo | CoB |
|---|---|---|---|---|
| Flower number | A↑\|D↓ | 13.56±0.60a | 14.67±0.53a | 14.15±0.77a |
| Leaf number | | 5.90±0.18 | 5.61±0.14 | 5.78±0.12 |
| Plant height (cm) | A↑\|D↓ | 30.43±0.59a | 29.39±0.40a | 25.19±0.56b |
| Flower diameter (mm) | A | 12.97±0.16a | 13.94±0.11b | 13.32±0.18a |
| Nectar (µl) | A | 1.45±0.13a | 1.32±0.11a | 1.55±0.16a |
| Total volatile emission | A | 6454.98±472.45a | 8440.33±607.64b | 8852.67±566.33b |
| Benzaldehyde | A | 3196.38±347.19a | 4697.29±511.2b | 4603.56±247.58b |
| 1-butene-4-ITC | D | 117.69±18.83a | 242.81±37.92b | 1596.53±175.88c |
| Methyl benzoate | D | 144.67±15.69a | 117.01±8.53a | 125.57±15.89a |
| Phenylethyl alcohol | D | 1.02±0.18a | 1.74±0.98b | 2.07±0.53b |
| 2-Amino benzaldehyde | | 1644.32±154.67a | 1550.74±147.72a | 1043.4±165.01b |
| p-Anisaldehyde | A | 18.41±4.60a | 31.95±3.89b | 13.78±2.51a |
| Methyl anthranilate | D | 150.38±17.06a | 131.26±11.72a | 240.87±43.24b |
| (*Z*)-3-Hexenyl acetate | D | 99.55±10.6a | 182.58±32.26b | 134.74±10.28ab |
| Phenylacetaldehyde | A | 203.06±45.76ab | 381.81±134.72a | 133.41±35.99b |
| Benzyl nitrile | | 148.91±22.55ab | 173.16±14.83a | 109.73±17.09b |
| Methyl salicylate | D | 135.69±15.26a | 204.49±29.31b | 111.15±10.10a |
| Indole | D | 245.29±32.06a | 341.4±29.28b | 275.49±31.98ab |
| (*Z,Z*)-α-Farnesene | A | 10.36±0.81a | 13.19±0.62b | 14.91±1.35b |
| (*E,E*)-α-Farnesene | A | 339.23±36.68a | 370.9±21.28a | 447.45±42.78a |

**Table S4.** Values (mean ± SE) for flower visitation, oviposition, and herbivore resistance of plants and butterflies after 6 generations of selection under different biotic treatments in the ambient environment. H = hand pollination, Coevo = coevolution, CoB = coevolution + bumblebees. Herbivore resistance is measured as caterpillar weight (mg). Letters indicate significant differences at P ≤ 0.05 based on Tukey's post hoc tests comparing biotic treatments within the ambient environment. Each treatment consisted of 68-144 plants. Hypersensitive response (HR) as proportion of egg with necrotic (black) plant tissue under and around the egg (necrotic eggs / total eggs).

| | H | Coevo | CoB |
|---|---|---|---|
| Flower visitation | 156.93 ± 12.89a | 104.72±8b | 131.63±18.69ab |
| Number of eggs received | 18 ± 1.26a | 18.13±1a | 12.49±1.19b |
| Caterpillar weight (mg) | 136.01 ± 4.01a | 82.83±2.87b | 29.52±2.10d |
| Hypersensitive response | 0.05 ± 0.04a | 0.11 ± 0.01b | 0.79 ± 0.07c |

**Table S5.** Trait and fitness values (mean ± SE) of evolved butterfly traits after 6 generations of selection under different biotic treatments in the ambient environment. R = control (stock rearing), Coevo = coevolution, CoB = coevolution + bumblebees. Letters indicate significant differences at P ≤ 0.05 based on Tukey's post hoc tests comparing biotic treatments within the ambient environment. Each treatment consisted of 36-100 butterflies. Relative club length is the length of the club in proportion of total antenna length (club length / antenna length). Flower visitation speed in flowers visited per hour.

| | R | Coevo | CoB |
|---|---|---|---|
| Weight (mg) | 55.20±1.02a | 48.76±0.99b | 51.81±0.82ab |
| Size (wing area $cm^2$) | 2.04±0.02a | 1.83±0.02b | 1.95±0.02a |
| Antenna length (mm) | 10.35±0.05a | 10.05±0.05b | 10.27±0.05a |
| Club length (mm) | 2.07±0.02a | 2.11±0.03a | 2.09±0.03a |
| Relative club length | 0.19±0.00a | 0.21±0.00b | 0.20±0.00ab |
| Tongue length (mm) | 9.52±0.06a | 9.21±0.07a | 9.50±0.08a |
| Flower visitation speed | 28.39±2.39a | 22.88±1.82a | 26.23±3.30a |
| Number of eggs laid | 38.18±3.34a | 38.96±2.74a | 24.97±2.27b |

**Table S6.** Values (mean ± SE) for plant reproduction after 6 generations of selection under different biotic treatments in the ambient environment. H = hand pollination, Coevo = coevolution, CoB = coevolution + bumblebees. Letters indicate significant differences at P ≤ 0.05 based on Tukey's post hoc tests comparing biotic treatments within the ambient environment. Each treatment consisted of 68-144 plants. Fruit set as proportion of flowers developed into fruits (number of fruits / number of flowers). Damaged fruits as proportion of fruits with signs of damage (damaged fruits / number of fruits).

| | H | Coevo | CoB |
|---|---|---|---|
| Fruit set | 0.85±0.03a | 0.79±0.02a | 0.76±0.04a |
| Number of fruits | 14.39±0.81a | 14.77±0.68a | 14.3±1.17a |
| Damaged fruits | 0.23±0.03a | 0.23±0.02a | 0.03±0.01b |
| Number of seeds | 225.99±16.5a | 215.06±11.45a | 184.72±15.59a |
| Seeds per fruit | 15.18±0.56a | 14.63±0.47a | 13.99±0.74a |

**Table S7.** Trait values (mean ± SE) of evolved plant traits after 6 generations of selection under different biotic treatments in the hot environment. H = hand pollination, Coevo = coevolution, CoB = coevolution + bumblebees. Letters indicate significant differences at P ≤ 0.05 based on Tukey's post hoc tests comparing biotic treatments within the hot environment. Z- and P-values indicate significant differences comparing the ambient and hot environment for specific biotic treatments, where bold values indicate P ≤ 0.05 and italic values P ≤ 0.1 based on Tukey's post hoc tests. Scent emission is expressed as pg/flower/l/h. Each treatment consisted of 68-144 plants. ITC = isothiocyanate.

| | Hot environment | | | Compared to ambient | | | | | |
|---|---|---|---|---|---|---|---|---|---|
| | H | Coevo | CoB | H | | Coevo | | CoB | |
| | | | | z | P | z | P | z | P |
| Flower number | 11.36±0.47a | 11.67±0.67a | 17.88±0.99b | 2.28 | **0.022** | 3.50 | **0.001** | -3.10 | **0.002** |
| Leaf number | 5.42±0.16 | 5.15±0.16 | 6.06±0.19 | - | - | - | - | - | - |
| Plant height (cm) | 29.19±0.46a | 22.26±0.53b | 29.19±0.54a | 1.61 | 0.107 | 10.68 | **<0.001** | -5.19 | **<0.001** |
| Flower diameter (mm) | 13.01±0.12a | 13.61±0.16b | 13.34±0.18ab | -0.15 | 0.878 | 1.70 | *0.090* | -0.06 | 0.951 |
| Nectar (µl) | 1.10±0.12a | 1.61±0.17b | 0.97±0.13a | 1.83 | *0.068* | -1.64 | 0.101 | 3.03 | **0.003** |
| Total volatile emission | 7666.55±653.22a | 12821.80±696.59b | 7053.32±611.24a | -1.81 | *0.071* | -5.19 | **<0.001** | 2.57 | **0.010** |
| Benzaldehyde | 2597.65±199.53a | 6535.48±406.78c | 4719.44±512.60b | 2.13 | **0.033** | -3.93 | **<0.001** | 0.03 | 0.973 |
| 1-butene-4-ITC | 210.81±51.27a | 1056.02±131.95b | 288.95±56.54a | -3.44 | **0.001** | -10.04 | **<0.001** | 10.08 | **<0.001** |
| Methyl benzoate | 132.78±19.49ab | 165.53±15.29a | 121.97±10.63b | 0.66 | 0.508 | -3.25 | **0.001** | 0.32 | 0.751 |
| Phenylethyl alcohol | 1.51±0.29b | 2.82±1.22a | 0.59±0.05c | -2.51 | **0.012** | -3.51 | **0.001** | 6.62 | **<0.001** |
| 2-Amino benzaldehyde | 2812.88±358.43a | 2518.94±335.50a | 816.59±114.14b | -2.88 | **0.004** | -3.01 | **0.003** | 1.31 | 0.189 |
| p-Anisaldehyde | 16.81±3.27a | 38.32±7.71b | 24.81±4.64ab | 0.14 | 0.891 | -0.93 | 0.352 | -2.61 | **0.009** |
| Methyl anthranilate | 186.28±19.57a | 257.91±31.18a | 92.96±11.04b | -1.25 | 0.213 | -4.61 | **<0.001** | 5.51 | **<0.001** |
| (*Z*)-3-Hexenyl acetate | 127.04±22.83a | 218.60±49.17b | 168.55±21.38ab | -1.84 | *0.066* | -1.73 | *0.083* | -1.45 | 0.147 |
| Phenylacetaldehyde | 418.66±110.70a | 365.09±90.06a | 104.71±27.51b | -3.17 | **0.002** | -0.17 | 0.867 | 0.97 | 0.332 |
| Benzyl nitrile | 214.72±28.98a | 276.96±37.96a | 83.74±9.04b | -1.95 | *0.051* | -2.90 | **0.004** | 1.44 | 0.150 |
| Methyl salicylate | 92.65±9.99a | 198.08±20.78b | 77.98±11.35a | 2.51 | **0.012** | 0.24 | 0.808 | 2.33 | **0.020** |
| Indole | 319.76±33.21a | 538.53±64.23b | 253.38±33.73a | -1.48 | 0.139 | -2.92 | **0.004** | 0.70 | 0.486 |
| (*Z,Z*)-α-Farnesene | 16.43±1.32a | 19.24±1.07a | 10.71±0.81b | -4.64 | **<0.001** | -4.47 | **<0.001** | 3.17 | **0.002** |
| (*E,E*)-α-Farnesene | 518.57±48.27a | 630.27±43.93a | 288.94±26.96b | -3.55 | **<0.001** | -5.19 | **<0.001** | 3.47 | **0.001** |

**Table S8.** Values (mean ± SE) for flower visitation, oviposition, and herbivore resistance of plants and butterflies after 6 generations of selection under different biotic treatments in the hot environment. H = hand pollination, Coevo = coevolution, CoB = coevolution + bumblebees. Letters indicate significant differences at P ≤ 0.05 based on Tukey's post hoc tests comparing biotic treatments within the hot environment. Z- and P-values indicate significant differences comparing the ambient and hot environment for specific biotic treatments, where bold values indicate P ≤ 0.05 and italic values P ≤ 0.1 based on Tukey's post hoc tests. Each treatment consisted of 68-144 plants. Hypersensitive response (HR) as proportion of eggs with necrotic (black) plant tissue under and around the egg (necrotic eggs / total eggs).

| | Hot environment | | | Compared to ambient | | | | | |
|---|---|---|---|---|---|---|---|---|---|
| | H | Coevo | CoB | H | | Coevo | | CoB | |
| | | | | z | P | z | z | P | z |
| Flower visitation | 121.38±9.12b | 180.58±17.59a | 81.82±12.38c | 1.45 | 0.148 | -4.13 | 1.45 | 0.148 | -4.13 |
| Number of eggs | 19.65±1.24a | 9.9±0.83b | 21.29±1.63a | -0.80 | 0.424 | 6.49 | -0.80 | 0.424 | 6.49 |
| Caterpillar weight (mg) | 130.36±3.54a | 67.04±4.04c | 57.89±2.15c | 0.85 | 0.394 | 4.88 | 0.85 | 0.394 | 4.88 |
| Hypersensitive response | 0.11 ± 0.02a | 0.14 ± 0.02a | 0.12 ± 0.01a | -2.71 | **0.007** | -2.13 | -2.71 | **0.007** | -2.13 |

**Table S9.** Trait and fitness values (mean ± SE) of evolved butterfly traits after 6 generations of selection under different biotic treatments in the hot environment. R = control (stock rearing), Coevo = coevolution, CoB = coevolution + bumblebees. Letters indicate significant differences at P ≤ 0.05 based on Tukey's post hoc tests comparing biotic treatments within the hot environment. Z- and P-values indicate significant differences comparing the ambient and hot environment for specific biotic treatments, where bold values indicate P ≤ 0.05 and italic values P ≤ 0.1 based on Tukey's post hoc tests. Each treatment consisted of 36-100 butterflies. Relative club length is the length of the club in proportion of total antenna length (club length / antenna length). Flower visitation speed in flowers visited per hour.

| | Hot environment | | | Compared to ambient | | | |
|---|---|---|---|---|---|---|---|
| | R | Coevo | CoB | Coevo | | CoB | |
| | | | | t | P | t | P |
| Weight (mg) | 55.20±1.02a | 43.79±1.4c | 49.83±1.02b | 3.44 | **0.001** | 3.44 | **0.001** |
| Size (wing area $cm^2$) | 2.04±0.02a | 1.86±0.04b | 1.90±0.02b | -0.96 | 0.339 | -0.96 | 0.339 |
| Antenna length (mm) | 10.35±0.05a | 10.22±0.1a | 10.09±0.05a | -1.91 | *0.058* | -1.91 | *0.058* |
| Club length (mm) | 2.07±0.02a | 2.22±0.04b | 2.30±0.03b | -2.29 | **0.023** | -2.29 | **0.023** |
| Relative club length | 0.19±0.00a | 0.22±0.00b | 0.23±0.00c | -1.41 | 0.161 | -1.41 | 0.161 |
| Tongue length (mm) | 9.52±0.06a | 9.12±0.11ab | 8.88±0.10b | 0.74 | 0.460 | 0.74 | 0.460 |
| Flower visitation speed | 28.39±2.39b | 36.42±3.6a | 17.13±1.77c | -3.81 | **<0.001** | -3.81 | **<0.001** |
| Number of eggs laid | 38.18±3.34a | 20.37±2.52b | 43.8±2.88a | 5.10 | **<0.001** | -3.94 | **<0.001** |

**Table S10.** Values (mean ± SE) for plant reproduction after 6 generations of selection under different biotic treatments in the hot environment. Letters indicate significant differences at P ≤ 0.05 based on Tukey's post hoc tests comparing biotic treatments within the hot environment. Z- and P-values indicate significant differences comparing the ambient and hot environment for specific biotic treatments, where bold values indicate P ≤ 0.05 and italic values P ≤ 0.1 based on Tukey's post hoc tests. Each treatment consisted of 68-144 plants. Fruit set as proportion of flowers developed into fruits (number of fruits / number of flowers). Damaged fruits as proportion of fruits with signs of damage (damaged fruits / number of fruits).

| | Hot environment | | | Compared to ambient | | | | | |
|---|---|---|---|---|---|---|---|---|---|
| | H | Coevo | CoB | H | | Coevo | | CoB | |
| | | | | z | P | z | P | z | P |
| Fruit set | 0.88±0.03a | 0.69±0.04b | 0.65±0.04b | -0.80 | 0.422 | 3.97 | **<0.001** | 0.79 | 0.429 |
| Number of fruits | 13.4±0.65a | 9.82±0.74c | 12±0.86b | -0.71 | 0.476 | 5.63 | **<0.001** | 2.44 | **0.015** |
| Damaged fruits | 0.2±0.03a | 0.2±0.03a | 0.32±0.03b | 0.55 | 0.583 | 0.92 | 0.360 | -8.52 | **<0.001** |
| Number of seeds | 211.01±12.75a | 168.25±15.45b | 125.86±11.15b | -1.08 | 0.280 | 4.05 | **<0.001** | 3.46 | **0.001** |
| Seeds per fruit | 15.58±0.56a | 16.44±0.78a | 10.19±0.59b | -1.21 | 0.229 | -0.95 | 0.344 | 3.69 | **<0.001** |

**Table S11.** Effect of temperature and biotic environment on plant trait evolution after 6 generations of selection. Bold values indicate results where P ≤ 0.05 based on (generalized) linear mixed models. Italic values indicate results where P ≤ 0.1. Scent emission is expressed as pg/flower/l/h. Each treatment consisted of 68-144 plants.

| | Temperature (T) | | | Biotic (B) | | | T*B | | |
|---|---|---|---|---|---|---|---|---|---|
| | $\chi^2$ | df | *P* | $\chi^2$ | df | *P* | $\chi^2$ | df | *P* |
| Flower number | 2.81 | 1 | *0.093* | 24.35 | 2 | **<0.001** | 24.25 | 2 | **<0.001** |
| Leaf number | 1.12 | 1 | 0.291 | 3.67 | 2 | 0.159 | 3.22 | 2 | 0.199 |
| Plant height (cm) | 23.00 | 1 | **<0.001** | 41.86 | 2 | **<0.001** | 120.62 | 2 | **<0.001** |
| Flower diameter (mm) | 0.91 | 1 | 0.339 | 32.74 | 2 | **<0.001** | 2.00 | 2 | 0.368 |
| Nectar (µl) | 2.18 | 1 | 0.140 | 1.37 | 2 | 0.504 | 13.02 | 2 | **0.001** |
| Total volatile emission | 8.29 | 1 | **0.004** | 35.78 | 2 | **<0.001** | 28.45 | 2 | **<0.001** |
| Benzaldehyde | 1.71 | 1 | 0.191 | 94.86 | 2 | **<0.001** | 18.24 | 2 | **<0.001** |
| 1-butene-4-isothiocyanate | 7.47 | 1 | **0.006** | 155.77 | 2 | **<0.001** | 206.77 | 2 | **<0.001** |
| Methyl benzoate | 2.29 | 1 | 0.130 | 1.18 | 2 | 0.555 | 8.78 | 2 | **0.012** |
| Phenylethyl alcohol | 0.00 | 1 | 0.974 | 22.70 | 2 | **<0.001** | 62.93 | 2 | **<0.001** |
| 2-Amino benzaldehyde | 7.63 | 1 | **0.006** | 49.82 | 2 | **<0.001** | 11.42 | 2 | **0.003** |
| p-Anisaldehyde | 3.70 | 1 | *0.054* | 28.79 | 2 | **<0.001** | 4.02 | 2 | 0.134 |
| Methyl anthranilate | 0.40 | 1 | 0.526 | 0.70 | 2 | 0.703 | 52.71 | 2 | **<0.001** |
| (*Z*)-3-Hexenyl acetate | 8.36 | 1 | **0.004** | 34.56 | 2 | **<0.001** | 0.09 | 2 | 0.955 |
| Phenylacetaldehyde | 1.69 | 1 | 0.194 | 63.02 | 2 | **<0.001** | 9.34 | 2 | **0.009** |
| Benzyl nitrile | 4.48 | 1 | **0.034** | 43.10 | 2 | **<0.001** | 9.80 | 2 | **0.007** |
| Methyl salicylate | 7.83 | 1 | **0.005** | 62.99 | 2 | **<0.001** | 3.93 | 2 | 0.140 |
| Indole | 5.25 | 1 | **0.022** | 19.85 | 2 | **<0.001** | 6.00 | 2 | **0.050** |
| (*Z,Z*)-α-Farnesene | 13.42 | 1 | **<0.001** | 11.34 | 2 | **0.003** | 38.38 | 2 | **<0.001** |
| (*E,E*)-α-Farnesene | 11.12 | 1 | **0.001** | 9.18 | 2 | **0.010** | 40.82 | 2 | **<0.001** |

**Table S12.** Effect of temperature and biotic environment on flower visitation, oviposition and herbivore resistance of plants and butterflies after 6 generations of selection under different temperature and biotic environments. Herbivore resistance is measured as caterpillar weight (mg). Bold values indicate results where P ≤ 0.05 based on (generalized) linear mixed models. Italic values indicate results where P ≤ 0.1. Each treatment consisted of 68-144 plants and 36-100 butterflies. Hypersensitive response (HR) as proportion of egg with necrotic (black) plant tissue under and around the egg (necrotic eggs / total eggs).

| | Temperature (T) | | | Pollinator (P) | | | T*P | | |
|---|---|---|---|---|---|---|---|---|---|
| | $\chi^2$ | df | *P* | $\chi^2$ | df | *P* | $\chi^2$ | df | *P* |
| Flower visitation | 0.02 | 1 | 0.879 | 7.87 | 2 | **0.020** | 28.77 | 2 | **0.001** |
| Number of eggs | 0.58 | 1 | 0.447 | 11.68 | 2 | **0.003** | 68.32 | 2 | **<0.001** |
| Caterpillar weight (mg) | 7.09 | 1 | **0.008** | 766.99 | 2 | **<0.001** | 160.02 | 2 | **<0.001** |
| Hypersensitive response | 33.63 | 1 | **<0.001** | 150.63 | 2 | **<0.001** | 169.04 | 2 | **<0.001** |

**Table S13.** Effect of temperature and biotic environment on butterfly trait evolution and fitness after 6 generations of selection. Bold values indicate results where P ≤ 0.05 based on (generalized) linear mixed models. Italic values indicate results where P ≤ 0.1. Each treatment consisted of 36-100 butterflies. Relative club length is the length of the club in proportion of total antenna length (club length / antenna length). Flower visitation speed in flowers visited per hour.

| | Temperature (T) | | | Pollinator (P) | | | T*P | | |
|---|---|---|---|---|---|---|---|---|---|
| | $\chi^2$ | df | *P* | $\chi^2$ | df | *P* | $\chi^2$ | df | *P* |
| Weight (mg) | 11.55 | 1 | **0.001** | 15.41 | 1 | **<0.001** | 1.79 | 1 | 0.180 |
| Size (wing area $cm^2$) | 0.04 | 1 | 0.851 | 13.08 | 1 | **<0.001** | 2.81 | 1 | *0.094* |
| Antenna length (mm) | 0.11 | 1 | 0.746 | 1.08 | 1 | 0.300 | 6.49 | 1 | **0.011** |
| Club length (mm) | 18.58 | 1 | **<0.001** | 0.15 | 1 | 0.695 | 2.74 | 1 | *0.098* |
| Relative club length | 18.99 | 1 | **<0.001** | 0.00 | 1 | 0.979 | 8.41 | 1 | **0.004** |
| Tongue length (mm) | 11.62 | 1 | **0.001** | 0.38 | 1 | 0.537 | 7.78 | 1 | **0.005** |
| Flower visitation speed | 1.88 | 1 | 0.170 | 5.57 | 1 | **0.018** | 17.65 | 1 | **<0.001** |
| Number of eggs laid | 1.42 | 1 | 0.233 | 0.58 | 1 | 0.445 | 40.12 | 1 | **<0.001** |

**Table S14.** Effect of various factors on the reproduction of plants after 6 generations of selection under different temperature and biotic environments. Bold values indicate results where P ≤ 0.05 based on (generalized) linear mixed models. Italic values indicate results where P ≤ 0.1. Each treatment consisted of 68-144 plants.

| | | | | **Plant** | **Fruit set** | | | |
|---|---|---|---|---|---|---|---|---|
| Temperature (T) | | | | | | $\chi^2$ | df | *P* |
| Pollinator (P) | | | | | T | 7.07 | 1 | **0.008** |
| Number of visits (V) | | | | | P | 12.87 | 2 | **0.002** |
| Herbivory (H) | | | | | V | 56.75 | 1 | **<0.001** |
| Number of flowers (F) | | | | | H | 44.14 | 1 | **<0.001** |
| | | | | | - | - | - | - |
| **Number of fruits** | | | | | T*P | 9.90 | 2 | **0.007** |
| | $\chi^2$ | df | *P* | | **Damaged fruits** | | | |
| T | 20.49 | 1 | **0.006** | | T | 7.85 | 1 | **0.005** |
| P | 7.25 | 2 | **0.027** | | P | 1.44 | 2 | 0.487 |
| V | 58.87 | 1 | **<0.001** | | V | 0.35 | 1 | 0.554 |
| H | 57.20 | 1 | **<0.001** | | H | 123.94 | 1 | **<0.001** |
| F | 67.14 | 1 | **<0.001** | | F | 0.01 | 1 | 0.924 |
| T*P | 18.52 | 2 | *0.095* | | T*P | 65.29 | 2 | **<0.001** |
| **Number of seeds** | | | | | **Seeds per fruit** | | | |
| T | 15.37 | 1 | *0.088* | | T | 0.68 | 1 | 0.411 |
| P | 19.00 | 2 | *0.075* | | P | 28.31 | 2 | **0.001** |
| V | 79.99 | 1 | **<0.001** | | V | 48.50 | 1 | **<0.001** |
| H | 20.79 | 1 | **0.005** | | H | 0.68 | 1 | 0.409 |
| F | 37.51 | 1 | **<0.001** | | F | 2.61 | 1 | 0.107 |
| T*P | 15.02 | 2 | **0.001** | | T*P | 14.93 | 2 | **0.001** |
